# Deciphering the thiolactonization mechanism in thiolactomycin biosynthesis

**DOI:** 10.1101/2024.12.18.629141

**Authors:** Jiawei Guo, Qiaoyu Zhang, Yang Shen, Fangyuan Cheng, Moli Sang, Xuan Wang, Yunjun Pan, Mingyu Liu, Hao-Bing Yu, Bo Hu, Sheng Wang, Liangzhen Zheng, Ce Geng, Chaofan Yang, Lianzhong Luo, Gang Zhang, Lei Du, Yuanning Li, Wei Zhang, Yandong Zhang, Binju Wang, Shengying Li, Xingwang Zhang

## Abstract

Thiolactomycin (**1**), which features a unique *γ*-thiolactone ring, is a promising antibiotic candidate that specifically targets bacterial type II fatty acid synthase. Despite extensive studies on its pharmacological activities, modes of action, and chemical synthesis, the enzymatic processes responsible for forming the activity-determining *γ*-thiolactone ring have remained largely unknown. Here, we resolve this problem by revealing that the condensation and heterocyclization (Cy) domain of the nonribosomal peptide synthetase (NRPS) TlnC (TlnC_Cy_), along with the cytochrome P450 enzyme TlnA, cooperatively enable the *γ*-thiolactone assembly. TlnC_Cy_ mediates an unusual sulfurtransfer reaction to sulfurate the polyketide intermediate, generating a thiocarboxylate intermediate. Subsequently, TlnA acts as a *γ*-thiolactone synthase, converting the linear thiocarboxylate intermediate into **1** via a distal radical-based cyclization mechanism. These findings not only expand the functional and catalytic repertoires of NRPS Cy domains and P450 enzymes, but also highlight a special enzymatic strategy for *γ*-thiolactone biosynthesis in nature.

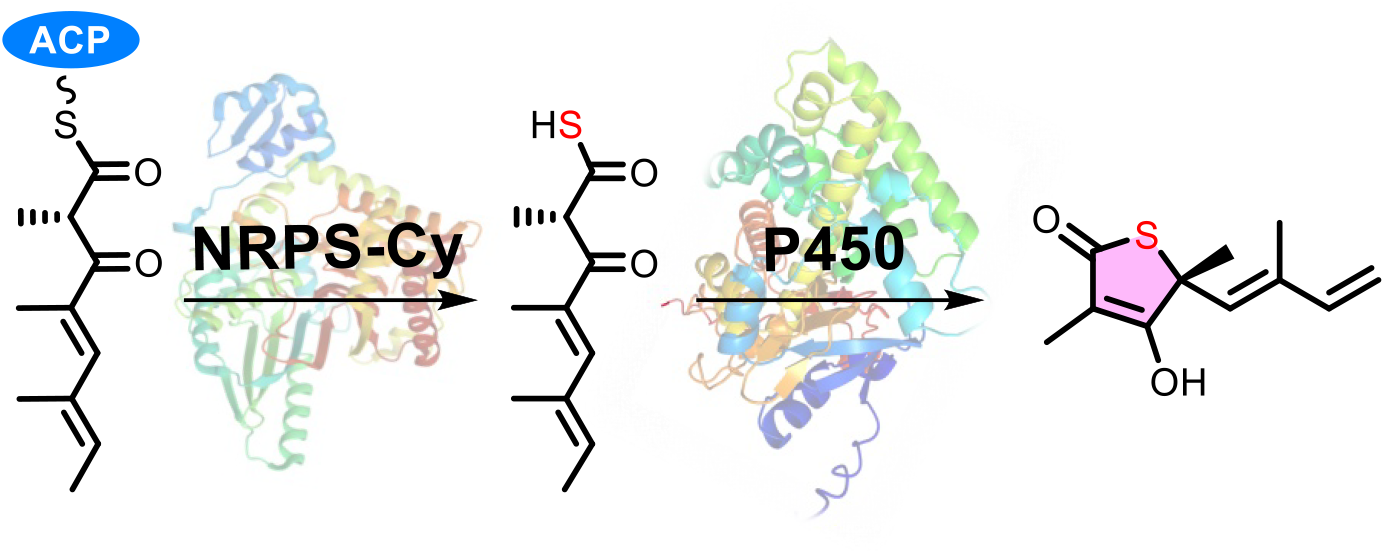

## 1 INTRODUCTION

Natural sulfur-containing molecules (SMs) have garnered significant attention in the field of biosynthesis, owing to their essential physiological and ecological functions, as well as their remarkable biological and pharmaceutical activities.^1–3^. Since the sulfur-bearing moieties are usually essential for the activities of SMs, the sulfur incorporation and tailoring processes during SM biosynthesis are of particular interest. Although biosynthetic mechanisms for assembling these sulfur-containing moieties have been continuously elucidated^4–9^, the identification of the sulfur origin and characterization of the sulfur-metabolizing enzymes and relevant catalytic mechanisms remain challenging^6–8^. This is primarily due to the complex reactivity of sulfur and limited understandings of the enzymatic and chemical diversity involved in sulfur metabolic processes^10^.

Thiolactomycin (**1**) and its analogues (**2**-**4**), which feature a unique *γ*-thiolactone ring (Figure 1a), were originally isolated from *Lentzea* sp. ATCC 31319 in 1982 ^11, 12^. Compound **1** acts as a specific inhibitor of the bacterial type II fatty acid synthase FabB/FabF by mimicking the structure and conformation of the native malonyl-ACP substrate within the active site (Figure S1)^13^. This inhibition results in effective inhibitory activity against both Gram-positive and Gram-negative bacteria, while being non-toxic to mammalian cells^14, 15^. Consequently, **1** has long been considered a novel antibiotic candidate, as no clinical antibiotics can target bacterial FabB/FabF to the best of our knowledge. Although the pharmacological activities, modes of action, and chemical synthesis of **1** have been extensively studied over the past four decades^16–18^, its biosynthesis remains partially understood, despite considerable research efforts^19–25^.

**Figure 1.**
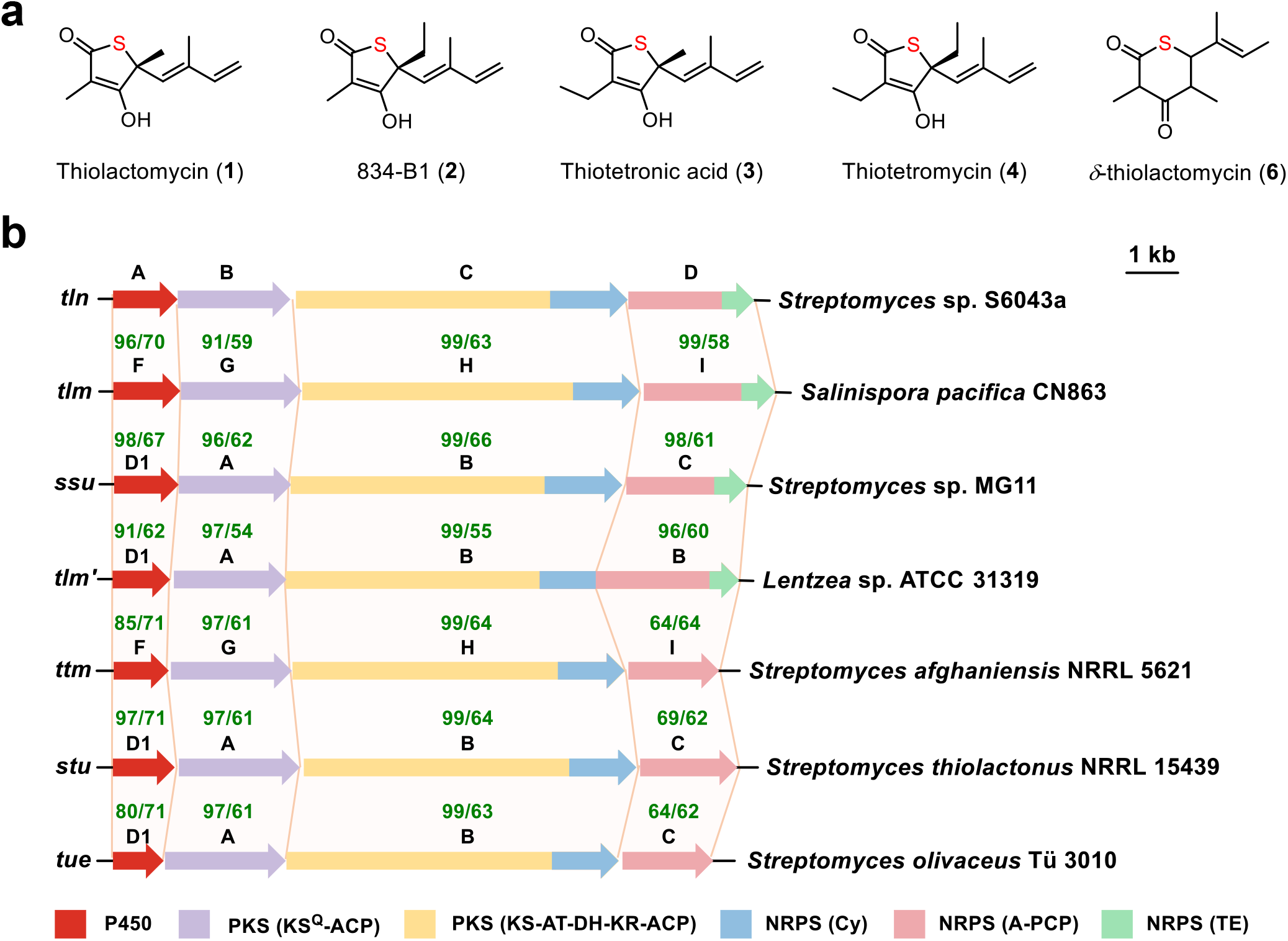
Chemical structures and biosynthetic gene clusters (BGCs) of thiolactomycin (1) and its analogues. **a**, Chemical structures of **1** and its analogues. **b**, Comparison of the BGCs of **1** from different microorganisms. The coverage/identity percentages (%/% in green) of the proteins encoded by the *tln* cluster with the corresponding homologues are shown above the gene names. Abbreviations: KS^Q^, malonyl-CoA decarboxylase; ACP, acyl carrier protein; KS, ketosynthase; AT, acyltransferase; DH, dehydratase; KR, ketoreductase; Cy, condensation and heterocyclization domain; A, adenylation domain; PCP, peptidyl carrier protein; TE, thioesterase.

The biosynthetic investigations for **1** were initiated by Reynolds and coworkers, and _L_-cysteine was identified as the origin of sulfur in this compound and its derivatives^19^. Then Moore and Leadlay/Sun groups identified the biosynthetic gene clusters (BGCs) and several key genes for the biosynthesis of **1**^20–23, 25^. Specifically, these studies have experimentally confirmed that the polyketide synthases (PKSs) encoded by the corresponding BGCs (Figure 1b) are responsible for building the carbon skeleton of **1** via consecutive decarboxylative condensations of a malonyl-CoA starter and three units of methylmalonyl-CoA extender (or random incorporation of one or two ethylmalonyl-CoA extender for producing **2**-**4**) with different levels of reduction, resulting in an acyl carrier protein (ACP)-tethered tetrapolyketide intermediate **5** (Figure 2a)^21–23^. Moreover, a cytochrome P450 enzyme was found to play an indispensable role in the construction of the *γ*-thiolactone ring. This conclusion was based on the observation that omission of the P450 encoding gene in the heterologous expression strain abolished the production of **1** and instead led to the formation of a *δ*-thiolactone ring-bearing product **6** (*δ*-thiolactomycin, Figures 2a and S2)^25^. As a result, two distinct hypotheses have been proposed regarding the thiolactonization process. One study suggested that a genetic locus encoding a cysteine desulfurase (StuS) and a thiouridylase-like sulfurtransferase (StuJ), located outside the BGC, facilitates the transfer of the sulfur atom from _L_-Cys to StuJ and then to the polyketide chain, thereby forming the thiolactone ring in **1** via a P450-derived epoxy intermediate (Figure S2a)^21^. The other study proposed that the BGC-encoded nonribosomal peptide synthetase (NRPS) directly incorporates the sulfur atom from _L_-Cys into the linear tetraketide via an unknown enzymatic process, with the P450 enzyme subsequently converting the thiocarboxyl intermediate into **1** via an epoxy intermediate (Figure S2b)^25^. Despite these two distinct hypotheses, the essential mystery of how the cysteine-derived sulfur atom is enzymatically incorporated into the thiolactone ring and the catalytic mechanism of the thiolactone-forming P450 still remain unknown.

**Figure 2.**
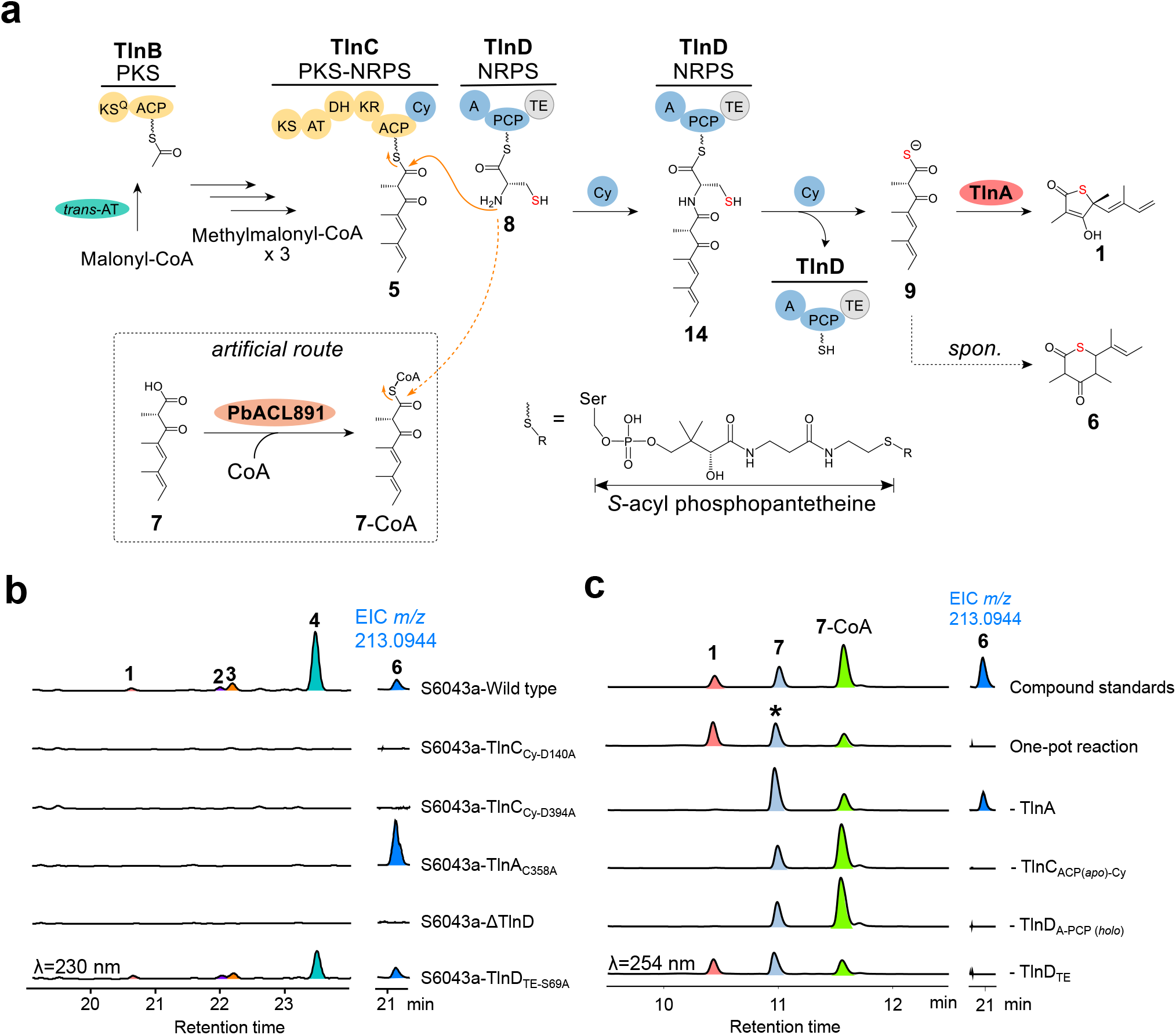
Elucidation of the thiolactomycin biosynthetic pathway. **a**, The elucidated biosynthetic pathway of **1**. The dashed boxed bypass shows the artificial route to generate **7**-CoA as a substrate mimic from synthetic **7**. The TE domain in TlnD is marked in gray since it is not required for **1** biosynthesis. Abbreviation: *trans*-AT, *trans*-type acyltransferase. **b**, HPLC (*left*) and HPLC-MS (*right*) analyses of production of **1**−**4** and **6** by *Streptomyces* sp. S6043 wild type and its mutant strains. **c**, HPLC (*left*) and HPLC-MS (*right*) analyses of the *in vitro* reconstituted one-pot reaction, along with control reactions where indicated enzymes were individually omitted. The one-pot reaction contained **7**-CoA, TlnC_ACP*(apo)*-Cy_, TlnD_A-PCP*(holo)*, L_-Cys, ATP, MgCl_2_, TlnD_TE_, TlnA, Fdx1499, FdR0978, NADPH, and DTT. (*: **7** was produced from hydrolysis of **7**-CoA; *see* Figure S16).

During our ongoing efforts to investigate diverse SM biosynthetic pathways^9, 26^, we re-isolated compounds **1**−**4** from *Streptomyces* sp. S6043a, a strain derived from an Antarctic deep-sea sponge. Using this strain, herein, we comprehensively characterized the biosynthetic process of the *γ*-thiolactone ring in **1**, through complementary approaches including bioinformatics analysis, *in vivo* gene inactivation, and *in vitro* enzymatic activity reconstitution. Notably, we demonstrated for the first time that the Cy domain of the NRPS TlnC mediates an unusual carboxyl-sulfuration reaction, forming of a key thiocarboxylate intermediate. Furthermore, we identified the P450 enzyme TlnA as a *γ*-thiolactone synthase that converts the linear thiocarboxylate intermediate to **1** through a distal radical-based cyclization mechanism. The key mechanisms for the sulfuration and thiolactonization steps were elucidated by a combination of biochemical, synthetic, and computational approaches.

## 2. RESULTS AND DISCUSSION

### 2.1 Isolation and BGC Identification of 1 from *Streptomyces* sp. S6043a

*Streptomyces* sp. S6043a was previously isolated from an Antarctic deep-sea sponge sample^27^. During our ongoing searches for natural SMs, we re-isolated and purified **1**−**4** from the Gauze’s Synthetic Medium No.1 (supplemented with 3.4% sea salts) fermentation cultures (10 L) of this strain (Figures 2b and S3). The structures of these compounds were elucidated by nuclear magnetic resonance (NMR) and high-resolution electro-spray ionization-mass spectrometry (HRESI-MS) analyses (Figures S4-S9), which are well consistent with previously reported data^20^. In addition to compounds **1**−**4**, HPLC-HRESI-MS analysis of the fermentation extract of *Streptomyces* sp. S6043a also detected the production of compound **6**, which bears a *δ*-thiolactone ring, albeit in very low yield (Figures 2b, S3 and S4).

The 6.9 Mbp whole genome of *Streptomyces* sp. S6043a was sequenced, assembled, and subsequently analyzed using antiSMASH 6.0^28^ to predict potential BGCs (Table S1). To identify the BGC of **1**−**4**, we used the known skeleton-building enzyme TlmH^20^ as a probe to scan the genome. Bioinformatics analysis of the candidate BGC (*tln*; GenBank accession number: OR392570) revealed the presence of all necessary genes for **1** biosynthesis, including the P450 gene *tlnA*, PKS gene *tlnB*, PKS-NRPS hybrid gene *tlnC*, and NRPS gene *tlnD* (Figure 1b). The four proteins encoded by these genes exhibit high sequence similarity to their corresponding homologues identified in other strains that also produce compound **1** (Figure 1b)^20, 21^.

Of note, careful sequence analysis of TlnC revealed intriguing information about the predicted Cy domain. Cy domains belong to the NRPS condensation domain (C domain) family. Unlike a typical C domain that catalyzes the formation of a single peptide bond, a Cy domain usually serves dual functions. These include condensation of an amino acid (typically cysteine or serine) and heterocyclization, forming of a thiazoline or oxazoline moiety (Figure S10)^29^. Protein sequence alignment revealed that the Cy domain in TlnC (TlnC_Cy_) and its homologous proteins share a conserved condensation motif (DXXXXD) with other typical Cy domains. However, the essential T-D catalytic dyad, which is crucial for heterocyclization in typical Cy domains, is replaced by an N-D motif in TlnC and its close homologs. (Figure S11)^30–32^. This difference suggests that the Cy domain in TlnC may play an unconventional role in the biosynthesis of **1**.

### 2.2 New Insights Provided by Gene Inactivation Experiments

According to the previous studies, deletion of the PKS encoding gene *tlmA’*, PKS-NRPS hybrid gene *stuB*, and P450 gene *stuD1* in *Lentzea* sp. ATCC 31319 or *Streptomyces thiolactonus* NRRL 15439 (Figure 1b) resulted in the complete abolition of **1** production, and no accumulation of potential intermediates was observed^21^. However, in a heterologous expression strain of the *tlm* BGC in *Streptomyces coelicolor* M1152, the omission of the P450 gene *tlmF* led to the identification of compound **6** bearing a *δ*-thiolactone ring, which was deduced to be a by-product derived from a thiocarboxyl intermediate (Figure S2b)^25^. These results provide limited but inspiring information regarding the details of thiolactonization and the role of compound **6** in the biosynthesis of **1**. To gain further insights into the potential functions of these genes, we conducted more precise and selective gene inactivation, as well as mutagenesis analysis for these genes in *Streptomyces* sp. S6043a.

Although the deletion of the entire PKS-NRPS hybrid gene *stuB* (the counterpart of *tlnC*) in *S. thiolactonus* NRRL 15439 strain confirmed its indispensable role in **1** biosynthesis^21^, the specific function of the uncommon Cy domain has yet to be investigated. Thus, we performed CRISPR-Cas9-based site-directed mutagenesis^33, 34^ *in vivo* to individually target the two subdomains of TlnC_Cy_. In the condensation subdomain, we introduced a single mutation by replacing the catalytically essential D_140_ residue with alanine in the D_135_XXXXD_140_ motif (Figure S11), generating the mutant strain S6043a-TlnC_Cy-D140A_^35^. In the heterocyclization subdomain, the putative catalytic residue D_394_ of the N_352_-D_394_ dyad was mutated to alanine, resulting in the mutant strain S6043a-TlnC_Cy-D394A_^31^. Product analysis revealed that the production of **1**−**4** was completely abolished in both mutant strains (Figure 2b). These results not only indicate the catalytic necessity of these two aspartic acid residues, but also suggest that both subdomains in TlnC_Cy_ are essential for the biosynthesis of **1**.

Mutating the C_358_ residue in P450 TlnA, which is responsible for coordinating with the heme-iron catalytic ligand in the active site and is thus absolutely conserved in all P450 enzymes (Figure S12)^36^, to alanine also cancelled the production of **1**−**4** (Figure 2b). However, HPLC-HRESI-MS analysis of the fermentation extract of this mutant strain (S6043a-TlnA_C358A_) detected the accumulation of compound **6** (Figure 2b), consistent with the previous report by Tang *et al*. on heterologous expression experiments^25^. Notably, **6** was absent in the fermentation extracts of both S6043a-TlnC_Cy-D140A_ and S6043a-TlnC_Cy-D394A_ mutants (Figure 2b). These findings strongly suggest that TlnC_Cy_, rather than TlnA, is involved in the sulfur incorporation process. It is likely that TlnA should be responsible for converting a sulfur-bearing intermediate (**6** or an unknown intermediate) into **1**.

The in-frame deletion of the entire NRPS gene *tlnD* (S6043a-ΔTlnD, Figure S13) led to complete abolishment of the production of **1**−**4** and **6** (Figure 2b), indicating the essential role of this gene in the biosynthesis of **1**. Surprisingly, targeted disruption of the thioesterase (TE) domain of TlnD by replacing the catalytically essential S_69_ residue with alanine (Figure S14) did not influence the production of **1**−**4** and **6** in the resulting S6043a-TlnD_TE-S69A_ strain (Figure 2b). Sequence analysis further revealed that three homologous proteins to TlnD—StuC from *S. thiolactonus* NRRL 15439, TtmI from *S. afghaniensis* NRRL 5621, and TueC from *S. olivaceus* Tü 3010—also lack a TE domain (Figure 1b and S15). Interestingly, despite the absence of a TE domain, these strains are still capable of producing **1**^20,21^. Taken together, these findings indicate that the TE domain embedded within the NRPS is not required for the biosynthesis of **1**. However, this raises an important question on how the PKS-NRPS hybrid product is released from the carrier protein.

### 2.3 Reconstitution of Thiolactonization *In Vitro*

To understand the enzymatic basis and catalytic details of the thiolactonization process, we attempted to reconstitute the entire biosynthetic pathway of **1** *in vitro*. Despite our best efforts, we were unable to express the full-length multi-domain proteins TlnC and TlnD in their soluble forms. To overcome this challenge, we turned to separately express the functional domains of TlnC and TlnD, focusing on those involved in *γ*-thiolactone construction. As a result, we successfully expressed the *N*-terminal SUMO-tagged TlnC_ACP-Cy_ di-domain, *N*-terminal His_6_tagged TlnD_A-PCP_ di-domain, and *N*-terminal His_6_-tagged TlnD_TE_ monodomain as stand-alone soluble proteins in *Escherichia coli* BL21(DE3) (Figure S17). Notably, the expression of TlnC_Cy_ and TlnD_A_ without the associated carrier proteins resulted in insoluble proteins. In addition, to obtain the activated *holo*-form of the carrier proteins with a phosphopantetheine arm, TlnC_ACP-Cy_ and TlnD_A-PCP_ were also expressed in the engineered *E. coli* BAP1 strain^37^, which harbors the *sfp* gene from *Bacillus subtilis*^38^ to enable the conversion of ACP/PCP from the inactive *apo*-form into the active *holo*-form. HPLC-HRESI-MS analysis confirmed that the purified TlnC_ACP-Cy_ and TlnD_A-PCP_ were both in the expected *holo*-forms (Figure S18). Moreover, the purified *N*-terminal His_6_-tagged P450 TlnA was verified to be active, as evidenced by the characteristic 450 nm peak in the CO-bound reduced difference spectrum (Figures S17 and S19)^39^.

Due to the inability to reconstitute the full function of TlnC *in vitro*, its predicted product **7** was chemically synthesized (Figure 2a and Scheme S1)^23^. To load **7** onto the PKS assembling line, we enzymatically prepared its CoA-linked form (**7**-CoA) as a mimic of the ACP-bound intermediate **5**, by co-incubating **7** and CoASH in the presence of a promiscuous acyl-CoA ligase PbACL891 from *Penicillium brevicompactum* NRRL864^40^ (Figure 2a). The structure of purified **7**-CoA was confirmed by NMR analysis (Figures S20-25). To investigate the function of TlnD_A-PCP_, the *holo* form protein was reacted with _L_-Cys (the predicted substrate based on analysis of the specificity code of A domain, *see* Table S2), ATP, and MgCl_2_. HPLC-HRESI-MS analysis of the reaction mixture revealed the expected product, cysteinyl-*S*-TlnD_A-PCP_ (**8**), along with two additional side products compared to the negative control reaction lacking ATP (Figure S26). Based on their molecular masses, we inferred that the two unexpected products were TlnD_A-PCP_ linked to two and three units of _L_-Cys, respectively (Figure S26). This phenomenon was likely due to artificial loading events mediated by the A domain under non-downstream conditions, where the second and third _L_-Cys molecules could attach to the exposed thiol group of **8** (Figure S27). Supporting this, only a single product peak was observed when *S*-methylcysteine, a cysteine derivative with its thiol group protected, was used in the reaction with TlnD_A-PCP*(holo)*_ (Figure S28).

Next, we sought to reconstitute the thiolactonization reaction using **7**-CoA as the starting material in a one-pot reaction. Specifically, **7**-CoA was co-incubated with TlnC_ACP*(apo)*-Cy_, TlnD_A-PCP*(holo)*_, TlnD_TE_ and TlnA, along with the surrogate P450 redox partners Fdx1499 and FdR0978 derived from the cyanobacterium *Synechococcus elongatus* PCC 7942^41^, as well as the co-factors _L_-Cys, MgCl_2_, ATP, NADPH and DTT for 4 h at 30 oC (Figure 2c). As expected, substantial production of **1** was observed, as confirmed by comparing the HPLC retention time and HRESI-MS spectrum with the isolated natural standard of **1** (Figures 2c and S4). Omitting TlnC_ACP*(apo)*-Cy_, TlnD_A-PCP*(holo)*_, or TlnA, respectively, abolished the production of **1**, whereas the absence of TlnD_TE_ had no effect on its formation (Figure 2c). This observation aligns with the *in vivo* TE inactivation result (Figure 2b), affirming that TlnD_TE_ does not participate in the biosynthesis of **1**. Therefore, the release of the PKS-NRPS product from the assembling line could occur spontaneously or through a non-conventional chain-termination mechanism that does not involve TE-mediated hydrolysis or cyclization. Taken together, these results demonstrate that TlnC_ACP-Cy_, TlnD_A-PCP_, and TlnA are necessary and sufficient to convert **7**-CoA into the final product **1**. Notably, the *δ*-thiolactone product **6** was detected only in the *in vitro* reaction without P450 TlnA (Figure 2c), further supporting the role of TlnA as a *γ*-thiolactone synthase.

### 2.4 *γ*-Thiolactone Formation Catalyzed by TlnA via a Distal Radical-Based Cyclization Mechanism

Since both *in vivo* inactivation and *in vitro* absence of TlnA blocked the formation of **1** and resulted in the production of the *δ*-thiolactone product **6**, we initially hypothesized that TlnA might convert **6** into **1** through an intramolecular rearrangement. However, when synthetic **6** (*see* Supporting Information) was reacted with TlnA in the presence of Fdx1499, FdR0978 and NADPH, no product was observed (Figure S29). This result ruled out **6** as a substrate for TlnA and confirmed it as a by-product formed during the sulfur incorporation process, consistent with the proposal by the Moore group^25^. Another previously proposed possibility is that TlnA catalyzes an online reaction towards the TlnC tethered **7** (*i*.*e*., **5**, Figure S2b)^21^. To test this hypothesis, **7**-CoA was co-incubated with TlnC_ACP*(apo)*-Cy_ and the purified Sfp^38^ to facilitate the transfer of the **7**-linked phosphopantetheine arm onto Ser_108_ of TlnC_ACP*(apo)*-Cy_ to form TlnC_ACP-Cy_-**7** (Figures 2a and S30). After confirming the formation of TlnC_ACP-Cy_-**7** by HRESI-MS (Figure S30), we added TlnA along with its redox partners and NADPH to the enzymatically produced TlnC_ACP-Cy_-**7** *in situ*. However, no reaction was detected (Figure S31).

The Moore group previously hypothesized that **6** could be spontaneously generated through the nucleophilic attack on C5 by the terminal thiocarboxyl group (*i*.*e*., R-COS^-^) of the putative intermediate **9** (Figure S2b)^25^. To verify this hypothesis, we attempted to synthesize **9** by reacting **7**-CoA with (NH_4_)_2_S (ammonium sulfide) in order to cleave the thioester bond (Figure 3a)^42^. Interestingly, HPLC-HRESI-MS analysis of the reaction mixture revealed the direct formation of **6**, but not **9**, even under various reaction conditions using different concentrations of (NH_4_)_2_S (Figures 3b and S32). These results confirmed that **6** was likely derived from **9** via rapid spontaneous C−S bond formation, which might be too unstable to be detected by HPLC-HRESI-MS under the reaction conditions. Therefore, we turned to incubate the P450 TlnA system with the *in situ* generated **9** in the **7**-CoA and (NH_4_)_2_S reaction. As expected, **1** was produced when TlnA, redox partners, and NADPH were included in the reaction mixture of **7**-CoA and (NH_4_)_2_S (Figures 3b and S4). This indicated that **9** is the real substrate of TlnA, confirming the role of the P450 enzyme as a *γ*-thiolactone synthase.

**Figure 3.**
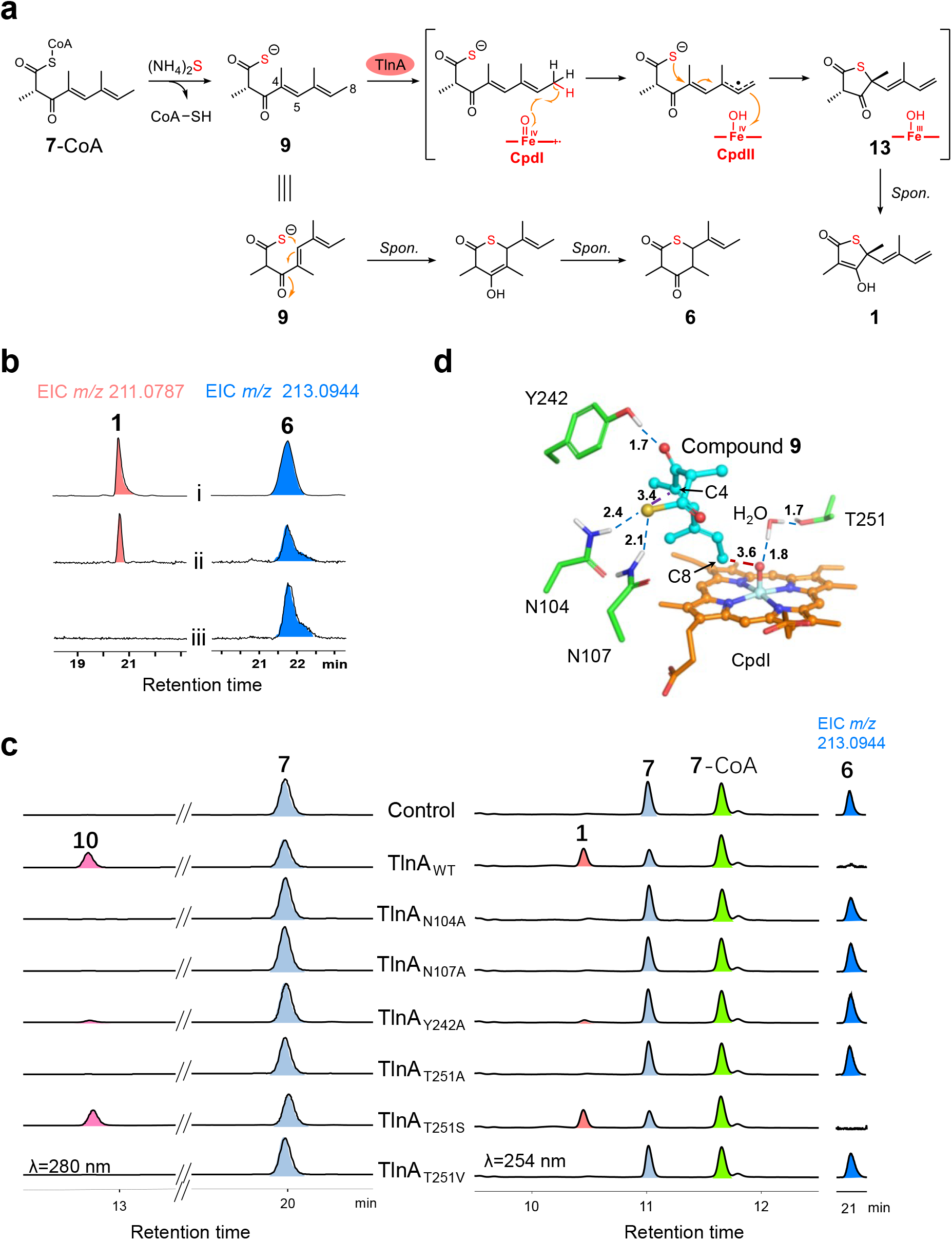
Functional and mechanistic analysis of P450 TlnA. **a**, Reaction scheme of the TlnA-mediated reaction. **b**, HPLC-MS analysis of the TlnA assays. (i) The authentic standards of **1** and **6**; (ii) The reaction of TlnA with **7**-CoA, Fdx1499, FdR0978, and NADPH in the presence of 20 mM (NH_4_)_2_S; (iii) the negative control of (ii) using boiling-inactivated TlnA. **c**, HPLC and HPLC-MS analysis of the catalytic activity of different TlnA mutants towards compound **7** (*left*) and the *in situ* generated **9** (*right*). The reaction system of **7** contained **7**, Fdx1499, FdR0978 and NADPH with TlnA wild type or its single mutants. The reaction system of **9** contained TlnC_ACP*(apo)*-Cy_, TlnD_A-PCP*(holo)*_, Fdx1499, FdR0978, **7**-CoA, and _L_-Cys with TlnA wild type or its single mutants in the presence of all necessary cofactors. Boiling inactivated wild-type TlnA (TlnA_WT_) was used in the negative control reactions. **d**, Molecular docking of **9** with TlnA. The red dashed line shows the distance between the C8 site of **9** and the oxygen atom of the CpdI in angstroms. The blue dashed lines indicate hydrogen bond interactions. The distance between the thiocarboxylic group and the C4 site of **9** is shown by the purple dashed line.

P450s are a superfamily of heme-thiolate enzymes that typically catalyze monooxygenation reactions in natural product biosynthesis^43–45^. Despite the ever-increasing examples of uncommon P450-catalyzed reactions (*e*.*g*., C−C, C−N, and C−S bond formations)^45–47^, to the best of our knowledge, TlnA represents the first example of a P450 enzyme catalyzing a *γ*-thiolactone formation reaction. In both previous hypotheses, TlnA was proposed to catalyze the epoxidation of the C4=C5 double bond in **5** or **9** to trigger the thiolactonization via bond rearrangement^21, 25^. To explore the catalytic mechanism of TlnA, we used the synthesized compound **7**, which is relatively stable in solution, as an analogue of substrate **9** to incubate with TlnA, its redox partners, and NADPH. As expected, TlnA catalyzed the transformation of **7** to a monooxygenated product **10** as detected by HPLC-HRESI-MS (Figures S4 and S33). However, due to the instability of **10** in the aqueous buffer, we were unable to obtain a sufficient amount of pure sample for direct structure determination, as it spontaneously decomposed to form **11** (Figures S4 and S33). Thus, we prepared the decomposition product **11** and determined its chemical structure by NMR analysis (Figures S34-S39). It was revealed that **11** should be the decarboxylation product of **10**. A similar decarboxylation reaction was also observed for **7**, spontaneously converting to **12** in the reaction buffer (Figure S33). Strikingly, the monooxygenation site of TlnA was determined to be the C8 methyl group

(for terminal hydroxylation), rather than the previously proposed C4=C5 double bond (for epoxidation). Given that **7** differs from **9** only by replacing the sulfur atom with an oxygen atom, we reasoned that the C8 methyl group in **9** should also be the oxidation site of TlnA during the native *γ*-thioesterification reaction. This is further supported by the primary kinetic isotope effect (KIE) experiment^48^, which showed that TlnA exhibited a significantly higher catalytic efficiency for the non-deuterated substrate **9** compared to the C8 deuterated substrate D_3_-**9** (*see* Supporting Information for details of chemical synthesis), with an approximately fivefold difference (*k*_H_/*k*_D_ = 5.59) (Figures S40-43). This KIE indicated that the rate-limiting C-H bond breaking indeed occurs at the C8 site.

According to the recognized P450 catalytic cycle^45^, we proposed two alternative mechanisms for TlnA. In the radical mechanism, following the binding of substrate **9** within the catalytic pocket of TlnA, the reactive species of heme-iron center (Compound I, CpdI) abstracts a hydrogen atom from the C8 methyl group, resulting in the formation of a substrate radical. Then, the nucleophilic thiocarboxylic group attacks the C4 position and initiates sequential double bond rearrangement and further electron transfer from the radical species to the ferrylhydroxo Compound II (CpdII), leading to the formation of the *γ*-thiolactone ring (Figure 3a). The generated Fe^III^-OH species coordinates with a proton from the solution to produce a H_2_O molecule and regenerates its initiating state to start a new catalytic cycle. In the alternative carbocation mechanism, after the formation of the substrate radical, CpdII of TlnA abstracts an electron from C8, producing a carbocation intermediate. Subsequently, the thiocarboxylic group attacks the C4 position and triggers double bond rearrangement to form the *γ*-thiolactone moiety (Figure S44). To distinguish these two mechanisms, we used ^18^O-labeled ^18^O_2_ or H_2_^18^O in the TlnA-**7** reaction to track the origin of the oxygen atom of the terminal hydroxyl group in **10**. HPLC-HRESI-MS analysis clearly indicated that the C8 oxygen atom of **10** should originate from ^18^O_2_ instead of H_2_^18^O (Figure S45). This observation strongly suggests that the TlnA-mediated hydroxylation of **7** likely undergoes the canonical ‘oxygen rebound’ process via the radical intermediate, but not through a carbocation intermediate, because otherwise the H_2_^18^O-derived ^18^O would be incorporated into **10** in the labelling experiments. Collectively, we reason that TlnA might adopt a similar radical mechanism when catalyzing its natural *γ*-thiolactone formation reaction.

Then, we attempted to use structural biology to gain more catalytic insights into TlnA. Due to unsuccessful attempts to crystallize TlnA, its 3D structure was predicted by AlphaFold3^49^ (Figure S46). Subsequently, **9** was docked into the active site of TlnA using AutoDock Vina^50^. The docked structure was then refined and optimized through molecular dynamics (MD) simulations (Figures 3d, 4, S47 and S48). The optimized complex showed that the distance between C8 of **9** and the ferryl-oxo group of CpdI is 3.6 Å (Figure 3d), while the distance from the closest hydrogen atom of C8-H_3_ to the oxygen atom of CpdI is 2.6 Å (Figure 4b). Such proximities are very suitable for the hydrogen atom transfer (HAT) from the substrate C8–H to CpdI (Figure 3d)^51^. This finding further supports that C8–H, rather than the C4=C5 double bond^21, 25^, should be selectively oxidized by TlnA.

**Figure 4.**
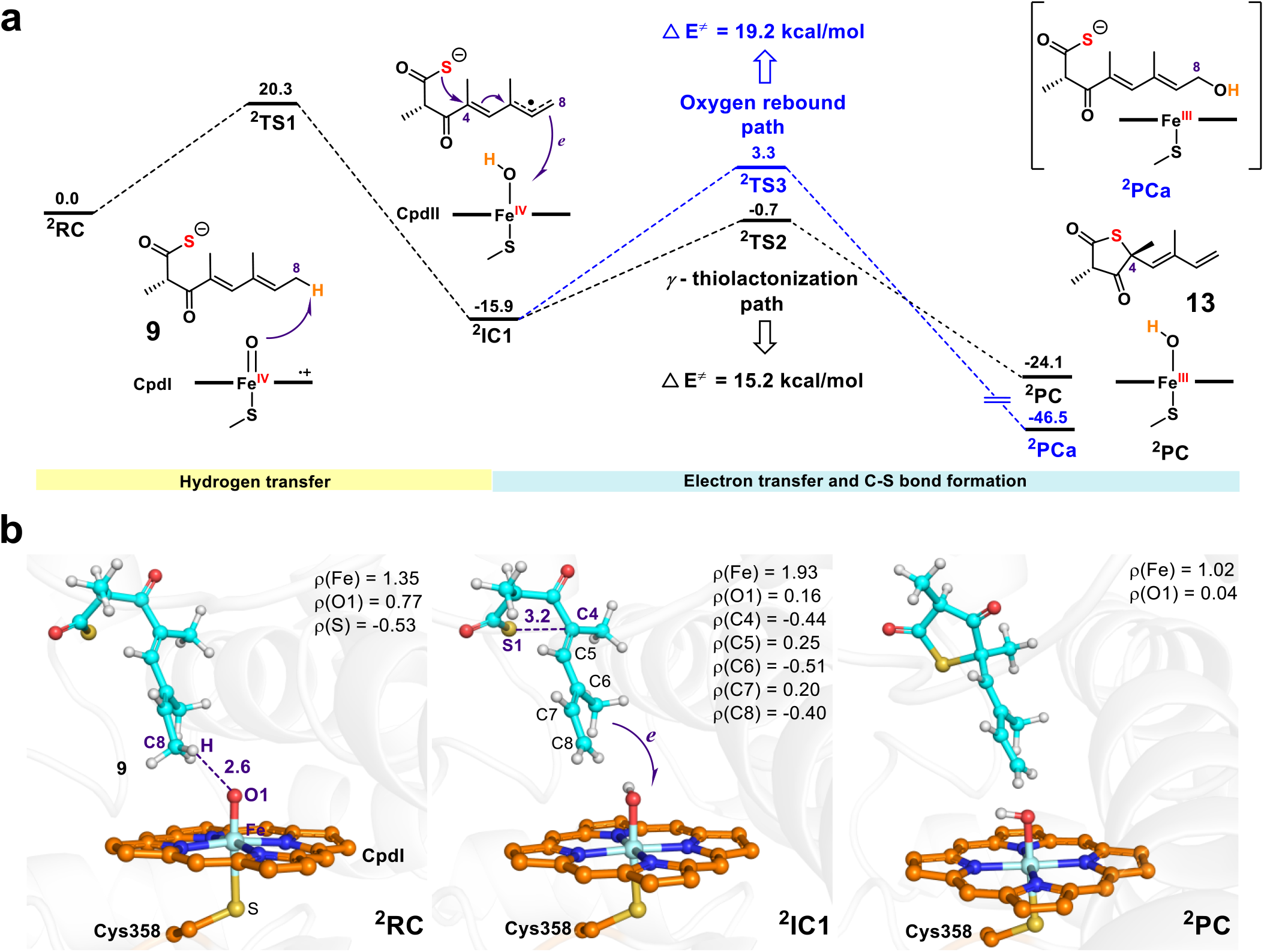
The calculated mechanism for the P450 TlnA-catalyzed oxidation of 9. **a**, QM(UB3LYP-D3BJ/B2)/MM-calculated energy profile (kcal/mol) for the P450 TlnA-catalyzed oxidation of **9** to generate **1. b**, QM(UB3LYP-D3BJ/B1)/MM-optimized structures of key species, shown along with the spin population of the key atoms. The distances are given in angstroms.

Furthermore, we verified the radical mechanism of TlnA using quantum mechanical/molecular mechanical (QM/MM) calculations (Figure 4). According to the calculation results, the HAT from the C8–H of **9** to CpdI overcomes a barrier of 20.3 kcal/mol, giving rise to the intermediate ^2^IC1, which is 15.9 kcal/mol lower in energy than the starting state ^2^RC. Population analysis revealed that the radical is delocalized across the conjugated chains in ^2^IC1, with significant populations at C8 (−0.40), C6 (−0.51) and C4 (−0.44). Additionally, to rule out the epoxide-based mechanism, we calculated the C4=C5 epoxidation process starting from ^2^RC (Figure S49). MD simulation and QM/MM calculations show that the distances between the oxygen atom of Fe(IV)=O and the C4 and C5 atoms of **9** are 8.2 Å and 7.1 Å, respectively, both of which are significantly longer than the distance between the C8 atom and Fe(IV)=O (3.6 Å) (Figure 4b and Figure S49). Furthermore, the QM/MM calculations reveal that the epoxidation of the C4=C5 double bond involves an energy barrier of 45.0 kcal/mol (^2^RC→^2^TS4 in Figure S50), which is significantly higher than that of the C8 HAT reaction (20.3 kcal/mol in Figure 4a). Therefore, these results collectively support the notion that Fe(IV)=O initiates a HAT reaction at the C8 position rather than promoting the epoxidation of the C4=C5 double bond.

In ^2^IC1 for the HAT pathway, the sulfur atom (S1) was observed to maintain a short distance of 3.2 Å from the C4 position in **9**, suggesting that the intramolecular nucleophilic attack of sulfur on C4 in the catalytic pocket is feasible. QM/MM calculations showed that this nucleophilic attack step involves a barrier of 15.2 kcal/mol (^2^IC1→^2^TS2), resulting in the formation of the *γ*-thiolactone moiety in ^2^PC, which is 8.2 kcal/mol lower in energy than ^2^IC1. Notably, this intramolecular nucleophilic attack is accompanied by electron transfer from the substrate to the Fe^IV^-OH species (CpdII), forming of Fe^III^-OH in ^2^PC (Figure 4a). The Fe^III^-OH then restores to the initiating state Fe^III^ by producing a H_2_O molecule. The keto-type product **13** in ^2^PC undergoes a keto-enol transformation, forming of the final product **1** (Figure S51).

Starting from ^2^IC1, we also calculated the energy barrier for the alternative ‘oxygen rebound’ route leading to the C8-hydroxylated product (^2^PCa). However, this pathway needs to overcome a high barrier of 19.2 kcal/mol (^2^IC1→^2^TS3 in Figure 4a). This finding well aligns with our experimental result that the P450 TlnA-catalyzed oxidation of **9** exclusively yielded **1**, and explains why the C8-hydroxylated product was not detected. To further understand the catalytic mechanism of TlnA, we investigated the TlnA-catalyzed hydroxylation process of **7** using MD simulations and QM/MM calculations (Figures S52-S54). Our simulations indicated that **7** adopts a similar binding conformation with **9** (Figure S53). As a result, the C8 methyl group in **7** is positioned to the oxygen of Cpd I with a distance of 2.3 Å, leading to an energy barrier of 15.4 kcal/mol for the HAT from C8–H of **7** to Cpd I. Starting from the radical intermediate ^2^IC1’, we found that the barrier for intramolecular C4-O bond formation (26.3 kcal/mol) is much higher than that of the OH-rebound pathway (11.5 kcal/mol), thus favoring the production of the C8-hydroxylated product (Figure S54). This result is also consistent with our experimental results (Figure S33).

To validate the docking results and further explore the substrate-binding properties of TlnA, alanine-scanning experiments were conducted on the amino acid residues that form hydrogen bonds with the substrate, as identified by the TlnA-**9** model. These residues include N_104_, N_107_, and Y_242_ (Figures 3d and S17). Compound **7** and the *in situ* generated **9**, produced by co-incubation of **7**-CoA, _L_-Cys, TlnC_ACP*(apo)*-Cy_ and TlnD_A-PCP*(holo)*_ in the presence of necessary cofactors, were used as substrates to test the catalytic activity of these TlnA mutants (TlnA_N104A_, TlnA_N107A_ and TlnA_Y242A_), respectively. Product analysis showed that all three mutants completely or significantly lost their catalytic activity against both **7** and **9** (Figure 3c), confirming the key roles of these residues in substrate binding and activity.

We proposed that the amide groups in N_104_ and N_107_ and the phenolic hydroxy group in Y_242_ might form three hydrogen bonds with the sulfur atom and C3 carbonyl group in **9**, thereby stabilizing the catalytic conformation of the substrate (Figure 3d). MD simulation showed that the distances between the N1 of N_107_ and S atom of **9** (d1) fluctuates around 3.0-3.5 Å, while the distance between the H3 of N_242_ and the O2 atom (d3) of substrate fluctuates around 1.7-2.0 Å, suggesting that both N_107_ and Y_242_ could maintain stable hydrogen bonds with **9** (Figure S55b and d). Although the N2 atom of N_104_ and the S atom of **9** (d2) fluctuate between 3.0 and 5.0 Å (mostly within 3.0 to 4.0 Å), suggesting that N_104_ might also form a hydrogen bond with **9** (Figure S55c). Thus, these results support the hypothesis that N_104_, N_107_, and Y_242_ are responsible for maintaining the catalytic conformation of the substrate by forming hydrogen bonds with intermediate **9**. Of note, circular dichroism (CD) measurements indicated that these mutations did not cause significant conformational changes (Figure S56).

Additionally, MD simulation results also showed that the T_251_ residue binds with a water molecule, which further interacts with the ferryloxo group in CpdI (Figures 3d and S53). Several previous studies have proposed that the conserved T_251_ residue plays an important role in oxygen activation, proton delivery, electron transfer, and positioning of the heme-ligand water (Figure S12)^52–56^. To investigate the role of T_251_ in TlnA for the unusual *γ*-thiolactoniazation functionality, we made three mutants including TlnA_T251A_, TlnA_T251S_, and TlnA_T251V_. Activity analysis showed that TlnA_T251A_ and TlnA_T251V_, both of which lack the hydroxyl group in the side chain of residue 251, were dead mutants towards **9** and **7**. In contrast, TlnA_T251S_, maintaining a hy-droxyl group at residue 251, retained the catalytic activity (Figure 3c). These results confirmed the important role of the hydroxyl group in T_251_ for maintaining the catalytic function of TlnA. Taken together, P450 TlnA mediates an unprecedented *γ*-thiolactoniazation towards **9** via a distal radical-based cyclization mechanism with the aid of N_104_, N_107_, Y_242_, and T_251_ residues.

### 2.5 Thiocarboxyl Intermediate Formation Mediated by the Cy Domain of TlnC

Based on the results that co-incubation of **7**-CoA, TlnC_ACP*(apo)*-Cy_, TlnD_A-PCP*(holo)*,_ and _L_-Cys led to the formation of **6** via the unstable intermediate **9** (Figure 2a), we deduced that **9** is the direct product off-loaded from the PKS-NRPS assembly line. During the reaction co-mediated by TlnC_ACP*(apo)*-Cy_ and TlnD_A-PCP*(holo)*_, the A domain of TlnD is responsible for loading _L_-Cys onto the assembly line, with the ACP domain of TlnC and the PCP domain of TlnD serving as carrier proteins for substrate/intermediate delivery (Figure 5a). Therefore, the Cy domain of TlnC (TlnC_Cy_) is likely responsible for mediating the sulfurtransfer from the TlnD_PCP_-bound _L_-Cys to the TlnC_ACP_-bound **7** (Figure 5a). To understand the catalytic process of TlnC_Cy_, we performed alanine scanning mutagenesis on the putative catalytic residues, including D_140_, N_352_, and D_394_ within the condensation motif D_135_XXXXD_140_ and the heterocyclization motif N_352_-D_394_ (Figures S11 and S17).

**Figure 5.**
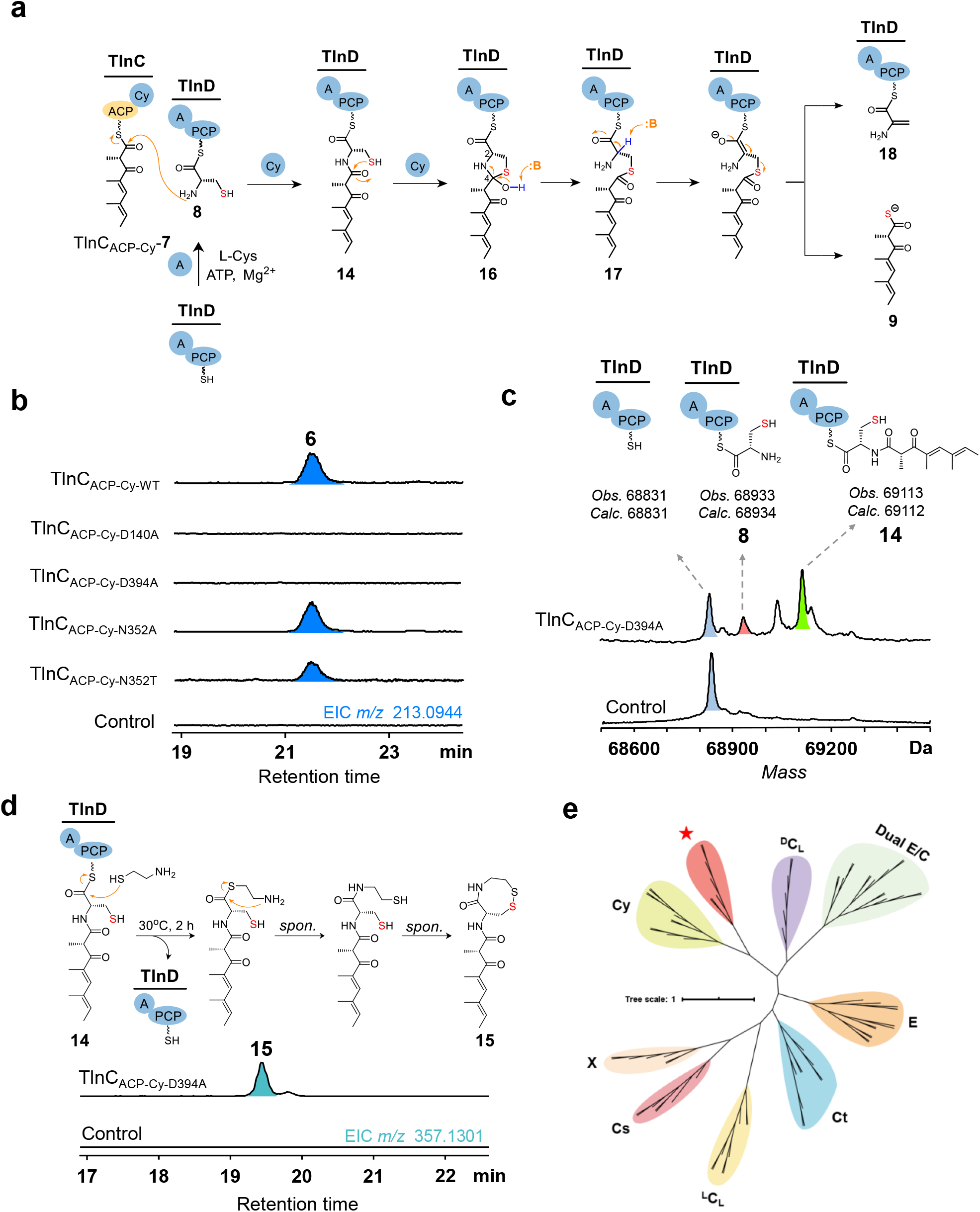
Function elucidation of the Cy domain of TlnC. **a**, Scheme of the reactions mediated by TlnCr_ACP*(apo)*-Cy_ and TlnDr_A-PCP*(holo)*_. **b**, HPLC-MS detection of the formation of **6** from the co-incubation of **7**-CoA, TlnDr_A-PCP*(holo)*_, r_L_-Cys, ATP, MgClr_2_ and DTT with TlnCr_ACP*(apo)*-Cy_ wild type or its single mutants, as well as the negative control using the boiling-inactivated TlnCr_ACP*(apo)*-Cy_. **c**, Deconvoluted ESI-MS analysis of the TlnDr_A-PCP_ bound intermediates resulted from the reaction of **7**-CoA, TlnCr_ACP*(apo)*-Cy-D394A_, TlnDr_A-PCP*(holo)*, L_-Cys, MgClr_2_, ATP and DTT; and ATP was not added in the negative control. **d**, Detection of the cysteamine captured TlnDr_A-PCP_ bound-intermediate. HPLC-MS analysis of the reaction of **7**-CoA with TlnDr_A-PCP*(holo)*_, TlnCr_ACP*(apo)*-Cy-D394A_ mutant, _L_-Cys, ATP, MgClr_2_ and DTT in the presence of 10 mM cysteamine; the negative control used the boiling-inactivated TlnCr_ACP*(apo)*-Cy-D394A_ mutant. **e**, Phylogenetic analysis of TlnCr_Cy_ and its homologous proteins. All the C domain sequences are collected from Conserved Domain Database (CDD)^32^. TlnCr_Cy_ and its homologous proteins (StuBr_Cy_, CUI25744.1; TueBr_Cy_, CUI25702.1; SsuBr_Cy_, CUI25727.1; TtmHr_Cy_, ALJ49921.1; TlmHr_Cy_, ALJ49910.1; TlmB’r_Cy_, CUI25672.1) are denoted by red star. Cy, condensation and heterocyclization; ^L^C_L_, condensation between two _L_-amino acids; ^D^C_L_, condensation between _D_- and _L_-amino acids; E, epimerization; Dual E/C, epimerization and condensation; Ct, fungal terminal C domain that terminates the macrocyclic product from NRPS via a TE-like hydrolytic mechanism; Cs, starting the NRPS assembly line; X domain, recruiting proteins to the assembly line.

Each single mutant (TlnC_ACP-Cy-D140A_, TlnC_ACP-Cy-D394A,_ or TlnC_ACP-Cy-N352A_) was individually incubated with TlnD_A-PCP*(holo)*_, **7**-CoA, and _L_-Cys in the presence of all necessary cofactors. Specifically, the TlnC_ACP-Cy-D140A_ and TlnC_ACP-Cy-D394A_ mutants both lost the ability to produce **6** (Figure 5b), which is consistent with the *in vivo* gene inactivation results (Figure 2b). In contrast, the TlnC_ACP-Cy-N352A_ mutant retained the catalytic activity (Figure 5b). These observations confirmed that both the condensation and heterocyclization catalytic motifs are necessary for TlnC to mediate the sulfurtransfer reaction, while the N_352_ residue, which is typically a threonine in Cy domains and crucial for the dehydration reaction forming the thiazoline moiety (Figure S10)^35^, does not participate in this reaction. Then, we further constructed the TlnC_ACP-Cy-N352T_ mutant to restore the T-D catalytic dyad of typical Cy domains and investigate whether this mutant could produce a thiazoline product. However, the N-to-T mutation neither affected the production of **6** nor resulted in the formation of any thiazoline product (Figures 5b and S57).

The mutation results suggest that the sulfurtransfer process likely includes two steps: the condensation of _L_-Cys and **7** and a heterocyclization-guided C−S bond cleavage reaction, with D_140_ and D_394_ serving as the catalytic residues. In principle, the TlnC_ACP-Cy-D394A_ mutant should retain the condensation activity but lose the ability to mediate the heterocyclization step, providing an opportunity to capture the online intermediate linked to TlnD_A-PCP_. Thus, we used HPLC-HRESI-MS to analyze the molecular weight changes of TlnD_A-PCP*(holo)*_ during the TlnC_ACP-Cy-D394A_-mediated transformation. As a result, the TlnD_A-PCP_-bound _L_-Cys (**8**) and the proposed condensation intermediate **14** were successfully detected (Figures 5a, 5c, and S57). To further confirm the structure of **14**, we used cysteamine as a chemical agent to trap it from TlnD_A-PCP_ (Figure 5d)^57^. When 10 mM of cysteamine was added to the reaction catalyzed by the TlnC_ACP-Cy-D394A_ mutant, a new product with an *m*/*z* value of 357.1313 ([M + H]^+^) was detected, which was identified as the cysteamine-trapped product **15** (Figures 5d and S4). These results confirmed **14** as the intermediate during the sulfurtransfer process (Figure 5).

After the formation of **14**, the sulfurtransfer process requires the cleavage of the C−S bond within the cysteine moiety (Figure 5a). Currently, pyridoxal 5’-phosphate (PLP) dependent cysteine desulfurases (CDS) and CDS-like enzymes are known as the main representatives for catalyzing C−S bond cleavage reactions^58, 59^. The mechanism of CDS/CDS-like enzymes involves sequential *α*-proton abstraction followed by *β*-elimination steps (Figure S58)^58, 59^. Intriguingly, during the assembly of the *β*-lactam skeleton of the nocardicin family of antibiotics, a special class of NRPS C domain was reported to employ a similar *α-* proton abstraction and *β*-elimination mechanism to mediate a C−O bond cleavage reaction towards the PCP-bound serine derivative substrate (Figure S59)^60–62^. Taken together, we proposed a putative mechanism for TlnCr_Cy_ to mediate the C-S bond cleavage reaction (Figure 5a). Initially, the two co-substrates, including the ACP-linked 7 and PCP-tethered L-Cys, bind to TlnCr_Cy_ to form a ternary complex. In the active site, the condensation subdomain first catalyzes the formation of **14**. Subsequently, the heterocyclization subdomain mediates an intramolecular cyclization to produce an unstable heterocyclic intermediate **16**, which then undergoes a spontaneous or Dr_394_-mediated C−N bond breakage reaction to form the linear intermediate **17**. Then, D_394_ or an unknown residue act as a general base, abstracting the *α*-proton in **17**^60–62^. This triggers the following *β*-elimination reaction, cleaving the C−S bond to produce **9** and the TlnDr_A-PCP_ tethered 2-aminoacrylic acid (**18**). 2-Aminoacrylic acid may be released from PCP spontaneously or via a residue-assisted hydrolysis reaction.

In a typical PLP-dependent CDS/CDS-like enzyme-mediated C−S bond cleavage reaction, the produced 2-aminoacrylic acid would spontaneously decompose into pyruvate and ammonia in solution (Figure S58)^63^. To examine whether pyruvate was formed in the TlnCr_Cy_-catalyzed reaction, 4-fluorophenylhydrazine (4-FPH) and *o*-phenylenediamine (OPD)^5^ were used as trapping agents for pyruvate^58^. HPLC-HRMS analyses of the 4-FPH and ODP treated reactions revealed a trace amount of pyruvate derivatives. However, comparable amounts of the pyruvate derivatives were also detected in the negative control reactions using boiled TlnCr_ACP-Cy_. This result suggested that the observed pyruvate was unlikely to be derived from the Cy catalytic process (Figure S60).

In principle, 2-aminoacrylic acid must be offloaded from TlnDr_A-PCP_, either enzymatically or spontaneously, or the catalytic cycle would be blocked. Supporting this, HR-ESIMS analysis of the molecular weight of TlnDr_A-PCP_ did not show any accumulation of **18**, with or without the inclusion of the TE domain of TlnD (Figure S61). Thus, we speculated that **18** might have gone through an unknown decomposition pathway to restore the unloaded TlnDr_A-PCP_ for the next round of catalysis. To investigate this hypothesis, we first attempted to detect potential volatile products using Gas Chromatography(GC), but no products were observed (Figure S62a). Then, we utilized dansulfonyl chloride (DC, to capture the potential NH_2_-containing products) and hydrazinopyridine (HP, to capture the potential carboxyl-containing products) to treat the TlnCr_ACP-Cy_ and TlnDr_A-PCP_ co-mediated one-pot reaction in order to trap any amino acid or carboxylic derivative resulting from **18**^64^. However, HPLC analysis showed that no new peak was observed in the DC/HP-treated samples compared to their corresponding control reactions using boiled TlnCr_ACP-Cy_ (Figure S62b and c).

Moreover, we attempted to use the ^13^C-tracer experiments to track the skeleton transformation of cysteine, using _L_-[1,2,3-^13^C_3_,^15^N] cysteine as the substrate. Initial attempts with TlnC_ACP-Cy_ and TlnD_A-PCP_ were unsuccessful due to their low catalytic activity. Therefore, we employed the homologous protein TlmB’, large PKS-NRPS hybrid protein derived from *Lentzea* sp. ATCC 31319 and showed much higher catalytic activity than TlnC_ACP-Cy_ and TlnD_A-PCP_ (Figures 1b, S17 and S63), for the ^13^C tracer experiments^23^. The reaction mixture containing _L_-[1,2,3-^13^C_3_,^15^N] cysteine, **7**-CoA, TlmB’, and essential cofactors in NaH_2_PO_4_ or Tris-HCl reaction buffer was monitored by time-course (1, 30, and 240 min) ^13^C-NMR spectroscopy. Notably, we observed the progressive conversion of cysteine-derived signals into two sets of new signals (Figure S64). Comparative spectral analysis revealed that while these new signals exhibited chemical shifts different from those of the substrate _L_-[1,2,3-^13^C_3_,^15^N] cysteine, their characteristic coupling patterns were remarkably conserved. This strongly suggests that the observed signals were likely derived from _L_-[1,2,3-^13^C_3_,^15^N] cysteine. The preserved coupling patterns suggest an intact C_3_ skeleton relative to _L_-[1,2,3-^13^C_3_,^15^N]cysteine, with the nitrogen atom still attached to C2 (as evidenced by the similar ^15^N-^13^C2 coupling constant of 3.6 Hz). The observed chemical shift variations indicate further decoration of the C_3_ skeleton during the *in vitro* reaction. Based on our proposed catalytic mechanism of the Cy domain, we proposed two possible derivatization pathways (Figure S65). Even though the current data are insufficient to determine the exact structure of the derivative of the C_3_ product, we are currently attempting to address this issue by employing a product release blockade strategy combined with structural biology approaches ^65^.

Next, we gauged the proposed mechanism using the cluster-continuum model calculations^66–68^, in which the methyl thioester compound **16’** was used as a simplified structural analog of **16** to reduce computational complexity (Figure S66). Starting from **16’**, water-assisted proton transfer from the hydroxyl group to the nitrogen atom generates a zwitterion intermediate. QM calculations showed that this step involves a barrier of 12.0 kcal/mol (RC′→TS1’). Then, a facile C−N bond cleavage leads to the ring-opened intermediate INT2’, which involves a slight barrier of 1.9 kcal/mol (INT1’→TS2’). Subsequently, a general base (D^394^ or another residue) abstracts the *α*-proton (INT2’→INT3’), initiating electron rearrangement and *β*-elimination, leading to the cleavage of the C−S bond and formation of **9** and **18’** (Figure S66). The overall process involves an energy barrier of 19.0 kcal/mol (INT2’→TS4’), giving rise to the final product **9** that is 8.5 kcal/mol lower in energy than **16’**, suggesting that this transformation is feasible both kinetically and thermodynamically.

To further explore the structural basis of the proposed mechanism for TlnC_Cy_, we used AlphaFold3 to predict the 3D structures of TlnC_Cy_ and TlnD_PCP_ (Figures S67 and S68). To reveal the structural similarities and differences between TlnC_Cy_ and typical Cy domains, the predicted structure of TlnC_Cy_ was aligned with several well-characterized conventional Cy crystal structures, including BmdB_Cy_, EpoB_Cy_, HMWP2_Cy_ and FmoA3_Cy_^31, 35, 69,70^. The results showed that TlnC_Cy_ exhibits a high degree of structural similarity with the classic Cy domains (with the root mean square deviation values ranging from 1.6 to 2.4 Å), both in overall structure and key catalytic residues (Figure S69 and S70). A detailed comparative analysis revealed that, near the catalytic center—particularly in close to the catalytic site D_394_—a loop and β-sheet formed moiety in TlnC_Cy_ is more extended than other Cy structures (Figure S69). This difference is also evident in the sequence alignment result (Figure S11). Based on these observations, we speculate that this unique structural feature may account for the difference in catalytic function and mechanism between TlnC_Cy_ and typical Cy domains. Moreover, TlnD_PCP_ tethered **14**, generated by AutoDock Vina, was docked to the active site of the TlnC_Cy_ model. We then refined and optimized the docking results by MD simulations (Figure S71). The refined complex model showed the typical catalytic conformation of Cy-PCP domains as that of BmdB_Cy2_-BmdB_PCP2_^35^. Particularly, it revealed a close distance of 3.1 Å between the D394 residue and the thiol group in **14** (Figure S72). This result supports the involvement of D_394_ in the transformation of intermediate **14**, consistent with our mutagenesis results (Figure 5b). However, unravelling the detailed catalytic mechanism requires a reliable complex crystal structure between TlnC_ACP-Cy_ and TlnD_A-PCP_, which is currently ongoing in our laboratories.

## 3. CONCLUSIONS

In this study, we elucidated the enzymatic basis and catalytic mechanisms underlying the *γ*-thiolactonization step in thiolactomycin biosynthesis. We demonstrated that the atypical Cy domain of TlnC mediates an unusual bond rearrangement to sulfurate the polyketide intermediate, producing the key thiocarboxylate intermediate **9**. Cladogram analysis reveals that TlnC_Cy_ and its homologs form a distinct clade separating from the typical Cy subfamily, suggesting their independent evolutionary pathway (Figure 5e). Recently, several uncommon Cy subfamily members have been found to catalyze various unusual reactions, including C−C bond formation and unconventional cyclizations^71–73^. Thus, the identification of TlnC_Cy_ as a thiocarboxy synthase further broadens the functional diversity of this subclass of enzymes and highlights their evolutionary divergence in natural product biosynthesis. Accordingly, this type of C domain could serve as a probe to mine more SMs that may deploy a TlnC_Cy_-like biosynthetic mechanism. It is worth noting that the thiocarboxylate group is rarely observed in natural products and is typically biosynthesized through a complicated sulfur incorporation process mediated by sulfur carrier proteins^74^. Therefore, our findings in this study present a new strategy for building the thiocarboxyl group in nature. However, despite employing multiple approaches to capture the predicted C_3_ byproduct released from intermediate **18**, its precise chemical structure remains unresolved. We are currently addressing this challenge by determining the complex structure of TlmB’ bound to the C_3_ product.

Furthermore, we determined the catalytic function of the P450 enzyme TlnA, which converts the thiocarboxylate intermediate **9** into the final product **1** by mediating a *γ*-thiolactone ring formation process. Considering the instability of **9**, we initially hypothesized that TlnA might interact with the NRPS assembly line, most likely via the Cy domain, to prevent **9** from converting into the side product **6** through substrate channeling effect. However, the measurements of potential TlnA-TlnC_ACP-Cy_ interaction by isothermal titration calorimetry (ITC) experiments, protein complex predictions by AlphaFold3, size-exclusion chromatography (SEC), and native-PAGE analyses all suggested that TlnA is unlikely to form a complex with the NRPS assembly line (Figures S73-S75). This may explain the formation of **6** from the spontaneous decomposition of **9** by the wild-type *Streptomyces* sp. S6043a.

In summary, we deciphered the long-standing *γ*-thiolactone biosynthetic mystery in thiolactomycin biosynthesis. Our results advance understanding of the functional diversity of both NRPS Cy domains and P450 enzymes, and underscore the strategic diversity for SM biosynthesis in nature. We envision that these findings will facilitate the future discovery, design, engineering, and application of these unusual enzymatic machineries.

## 4. METHODS

### 4.1 Materials

All antibiotics used in this study were obtained from Solarbio (Beijing, China). All amino acids and related derivatives were purchased from Heowns Biochem LLC (Tianjin, China). Organic solvents were acquired from Merck KGaA (Darmstadt, Germany). NADPH, ATP, DTT, and IPTG were bought from Aladdin (Shanghai, China). Cysteamine was purchased from McLean (Shanghai, China). 4-Fluorophenylhydrazine, dansyl chloride, and *o*-phenylenediamine were obtained from Solarbio (Beijing, China). Yeast extract and peptone were purchased from Angel Yeast Co.,Ltd (Yichang, China). The Ni-NTA Sefinose™ Resin (Settled Resin) for protein purification was acquired from Sangon Biotech (Shanghai, China). Thrombin for His-tag excision was acquired from Solarbio (Beijing, China). The ^18^O-labeled water was purchased from Tenglong Weibo (Qingdao, China). The isotopic ^18^O_2_ was obtained from Delin (Shanghai, China). The FlexiRun premixed gel solution for SDS-PAGE analysis was obtained from MDBio (Qingdao, China). Other chemicals and reagents were purchased from Sinopharm Chemical Reagent Co., Ltd. (Shanghai, China). Primers and other genes were synthesized by BGI (Shenzhen, China).

### 4.2 Software and tools

The BGCs of *Streptomyces* sp. S6043a were predicted using antiSMASH bacterial version^28^. NMR data was processed using MestReNova 9.0. All protein sequence alignments were performed on the T-Coffee website^75^. The NRPS Cy domains were aligned by MEGA11 program^76^ using Clustal W (version 11.0.13) with default parameters. The neighbor-joining phylogenetic tree was built with MEGA11 for 1200 bootstrap repetitions with default parameters^76^. The ESI-MS data were processed using Bruker Compass DataAnalysis version 4.2. Please refer to the Supporting Information for the software, tools, and detailed parameters used in the chemical calculations.

### 4.3 Analytical Methods

All HPLC analyses were conducted on a Thermo UltiMate 3000 instrument with a Triart C18 column (4.6 mm × 250 mm × 5 μM, YMC Co., Ltd., Japan). HPLC-HRESI-MS data of small molecules were recorded on a Bruker Maxis Q-TOF equipment with the same C18 column. HPLC-HRESI-MS analysis of proteins was performed on a Bruker Maxis Q-TOF equipment with a Triart bio C4 column (4.6 mm × 150 mm × 5 μM, YMC Co., Ltd., Japan). UV-vis absorbance spectra were taken using a SpectraMax M2 plate reader (Shanghai, China). NMR spectra were acquired on a Bruker Avance III 600 MHz spectrometer (Switzerland). DNA sequencing services were provided by Sangon Biotech (Shanghai, China). GC analyses were conducted utilizing an HP-5 column (30 m × 0.32 mm × 0.25 μm) on an Agilent Technologies 7890B instrument. The temperature program consisted of an initial hold at 80 °C for 1 minute, followed by a ramp to 220 °C at a rate of 5 °C/min, a subsequent hold at 220 °C for 1 minute, a further ramp to 290 °C at 3 °C/min, and a final hold at 290 °C for 15 minutes.

### 4.4 Growth conditions for production of 1 and its analogues by *Streptomyces* strains

*Streptomyces* sp. S6043a wild-type and mutant strains were inoculated on MS agar plates (20 g/L soybean flour, 20 g/L mannitol, and 20 g/L agar) and cultured at 30 °C for 72 h. A colony of each strain was then inoculated into 2×YT medium (16 g/L pancreatic peptone, 10 g/L yeast extract, and 5 g/L NaCl) and incubated for 36 h at 30 °C, 220 rpm. Then, the seed culture was inoculated into 50 mL of Gauze’s synthetic medium No.1 (20 g/L soluble starch, 1 g/L KNO_3_, 0.5 g/L K_2_HPO_4_, 0.5% g/L MgSO_4_·7H_2_O, 0.5 g/L NaCl, 0.01 g/L FeSO_4_·7H_2_O, 34 g/L sea salt, pH = 7.4) and further shaken at 30 °C, 220 rpm for 7 d to produce **1** and its analogues. The HPLC program for **1**−**4** analysis was as follow: 0 - 1 min, 10 % (ACN in H_2_O with 0.1% TFA); 2 - 25 min, 10 - 80 %; 26 - 30 min, 100 %; 30 - 35min, 10 %; 1 mL/min, 230 nm.

### 4.5 Isolation of 1 and its analogues

After cultivating *Streptomyces* sp. S6043a (10 L) for 7 days, sterilized XAD-16 resin (20 g/L) was added to absorb hydrophobic products. The culture-resin mixture was then shaken at 220 rpm for 2 h. The resin was filtered through cheesecloth, washed with deionized water, and eluted with acetone. The organic solvent was evaporated under reduced pressure, and the resulting aqueous layer was extracted with an equal volume of ethyl acetate (EtOAc) twice. The combined EtOAc fractions were dried *in vacuo* to yield a crude extract (10.6 g), which was then subjected to a silica gel column to separate **1** and its analogues from other components. The petroleum ether : EtOAc = 10 1 fraction was dried and subjected to HPLC for purification of **1** and its analogues using a YMC-Pack Pro C18 column (10 mm × 250 mm × 5 μM, YMC, Japan). The HPLC program was as follows: 0 - 1 min, 30 % (ACN in H2O); 2 - 20 min, 30 - 80 %; 20 - 25 min, 100 %; and 25 - 30 min, 30 %; 2.5 mL/min, 230 nm. The collected fractions were dried under vacuum to yield compounds **1−4** (**1**, 4.4 mg; **2**, 4.0 mg; **3**, 4.2 mg; **4**, 8.6 mg), the structures of which were determined by HRESI-MS and NMR analyses (*see* Supporting Information for details).

### 4.6 DNA isolation of *Streptomyces* sp. S6043a

The genomic DNA (gDNA) of *Streptomyces* sp. S6043a was extracted according to standard procedures^77^. The collected 2-d *Streptomyces* sp. S6043a mycelia were re-suspended in 5 mL of SET buffer (75 mM NaCl, 25 mM EDTA, 20 mM Tris, pH = 7.5). Then, lysozyme was added to the suspension with a final concentration of 1 mg/mL and incubated at 37 °C for 1 h. Then, 500 μL solution of 10% sodium dodecyl sulfate and 125 μL of 20 mg/mL proteinase K were added and co-incubated at 55 °C with occasional inversions for 2 h. After that, 2 mL of 5 M NaCl solution and 8 mL of phenol/chloroform/isoamyl alcohol (25:24:1) mixture were added to the clear solution and incubated at room temperature for 0.5 h with frequent inversions. The resulting suspension was centrifuged at 4,500 *×g* for 15 min, and the aqueous phase was transferred to a new tube using a blunt-ended pipet tip. An equal volume of isopropanol was then added to precipitate the total DNA, and the tube was inverted gently for 5 min before centrifugation at 4,500 *×g* for 5 min. The sedimentary DNA was transferred to a new microfuge tube and washed with 75% ethanol twice and dried under vacuum, until finally being dissolved in 1 mL of ddH_2_O containing 10 μg RNase.

### 4.7 Plasmids construction for CRISPR/Cas9 gene editing

All primers used in this study are listed in Table S2. The CRISPR/Cas9 gene editing plasmid pKCcas9dO^33^ was used as a template to generate the knockout and nucleotide site-specific mutagenesis plasmids. To construct nucleotide site-specific mutagenesis plasmids for TlnA, TlnC_D140A_, TlnC_D394A_ and TlnD_S69_, sgRNAs were designed at the mutation sites and amplified from pKCcas9dO as template using the primer pairs of TlnA-sgRNA-F/TlnA-sgRNA-R, TlnC_D140A_-sgRNA-F/TlnC_D140A_-sgRNA-R, TlnC_D394A_-sgRNA-F/TlnC_D394A_-sgRNA-R, and TlnD_S69_-sgRNA-F/TlnD_S69_-sgRNA-R, respectively. The homologous arm-pairs of the mutation sites for TlnA, TlnC_ACP-Cy-D140A_, TlnC_ACP-Cy-D394A_ and TlnD_TE-S69_ were amplified from the genomic DNA of *Streptomyces* sp. S6043a with the corresponding primer pairs of TlnA-up-F/TlnA-up-R and TlnA-dn-F/TlnA-dn-R, TlnC_ACP-Cy-D140A_-up-F/TlnC_ACP-Cy-D140A_-up-R and TlnC_ACP-Cy-D140A_-dn-F/TlnC_ACP-Cy-D140A_-dn-R, TlnC_ACP-Cy-D394A_-up-F/TlnC_ACP-Cy-D394A_-up-R and TlnC_ACP-Cy-D394A_-dn-FP/TlnC_ACP-Cy-D394A_-dn-RP, and TlnD_TE-S69_-up-F/TlnD_TE-S69_-up-R and TlnD_TE-S69_-dn-F/TlnD_TE-S69_-dn-R, respectively. To construct in-frame gene deletion plasmids for TlnD, the sgRNA was amplified from pKC-cas9dO as template with the primer pair of TlnDsgRNA-F/TlnDsgRNA-R. The homologous arm-pair of *tlnD* were amplified from gDNA of *Streptomyces* sp. S6043a as template with the corresponding primer pairs TlnD-up-F/TlnD-up-R and TlnD-dn-F/TlnD-dn-R. The sgRNAs and corresponding homologous arm-pairs were ligated into the *Spe*I*/Hind*III-digested pKCcas9dO using the ClonExpress Ultra One Step Cloning Kit and transformed into *E. coli* DH5α for plasmid amplification. The recombinant plasmids were verified by DNA sequencing and the correct plasmids were chemically transformed into *E. coli* ET12567. Finally, the plasmids were transformed into *Streptomyces* sp. S6043a by conjugation-transformation^78^ and the resultant recombinant strains were verified by DNA sequencing.

### 4.8 Plasmids construction for protein expression

The DNA sequence encoding TlnC_ACP-Cy_ was amplified from the gDNA of *Streptomyces* sp. S6043a using the primer pair of SUMO_ACP-Cy_-F/ SUMO_ACP-Cy_-R. The gene sequences of TlnA, TlnD_A-PCP_, TlnD_TE_, Sfp and TlmB’ were individually optimized and synthesized according to the codon preference of *E. coli*. The purified DNA fragments were ligated into the *Nde*I*/Hind*III-digested pET28a(+) or *Bam*HI*/Hind*III-digested pET28a-SUMO(+) using the ClonExpress Ultra One Step Cloning Kit, and transformed into *E. coli* DH5α for plasmid amplification. The recombinant plasmids were verified by DNA sequencing and the correct plasmids were chemically transformed into *E. coli* BL21 (DE3) for protein overexpression. Moreover, TlnC_ACP-Cy_, TlnD_A-PCP_ and TlmB’ were also expressed in *E. coli* BAP1 for converting ACP/PCP from the inactive *apo*-form into the active *holo*-form. The recombinant proteins of TlnD_A-PCP_, TlnD_TE_, Sfp and TlmB’ were produced in a version tagged with 6×His at their *N*-termini, while TlnA and TlnC_ACP-Cy_ were produced in a form tagged with SUMO and 6×His at their *N*-termini. The expression strains of *Sel*Fdx1499, *Sel*FdR0978 and PbACL891 were constructed in our previous studies_40, 41_.

### 4.9 Site-specific mutation of TlnA and TlnC_ACP-Cy_

The mutation sites of TlnA were introduced by site-directed PCR from pET28a-SUMO-TlnA as template using the primer pairs of TlnA_N104A_-F/TlnA_N104A_-R, TlnA_N107A_-F/TlnA_N107A_-R, TlnA_Y242A_-F/TlnA_Y242A_-R, TlnA_T251A_-F/TlnA_T251A_-R, TlnA_T251S_-F/TlnA_T251S_-R, and TlnA_T251V_-F/TlnA_T251V_-R (Table S2). The mutation sites of TlnC_ACP-Cy_ were introduced by site-directed PCR from pET28a-SUMO_ACP-Cy_ as template using the primer pairs of TlnC-ACP-Cy_D140A_-F/TlnC-ACP-Cy_D140A_-R, TlnC-ACP-Cy_D394A_-F/TlnC-ACP-Cy_D394A_-R, TlnC-ACP-Cy_N352A_-F/TlnC-ACP-Cy_N352A_-R, and TlnC-ACP-Cy_N352T_-F/TlnC-ACP-Cy_N352T_-R (Table S2). The amplified products were digested with *Dpn*I and purified using the Omega Gel Extraction Kit by following the manufacturers’ instructions and then transformed into *E. coli* DH5α for plasmid amplification. The recombinant plasmids were verified by DNA sequencing, and the correct plasmids were chemically transformed into *E. coli* BL21 (DE3) for protein expression.

### 4.10 Protein expression and purification

A single colony of each protein expression strain was inoculated into 5 mL LB broth containing kanamycin (50 mg/L) and grown at 37 °C, 220 rpm for 12 h. The resulting seed culture was inoculated into 0.5 L LB medium (supplemented with 4% glycerol) containing kanamycin (50 mg/L) in a 2 L conical flask, which was shaken and incubated at 37°C, 220 rpm for 4~6 h until the OD_600_ reached 0.6-0.8. Then, 0.2 mM IPTG was added to induce protein expression. For P450 TlnA, 0.25 mM 5-aminolevulinic acid (5-Ala) was further added as the heme precursor. The cells were further grown for an additional 16~20 h at 16 °C, 160 rpm, and then harvested by centrifugation at 6,000 *×g* for 10 min at 4 °C. The collected cells were suspended in 50 mL lysis buffer (50 mM NaH_2_PO_4_, 300 mM NaCl, 10 mM imidazole, 10% glycerol, pH = 8.0) and crushed by a high-pressure homogenizer (ATS, Shanghai, China) or an ultrasonic homogenizer JY92-IIDN (Scientz, Ningbo, China) at 4 °C. Bacterial cell debris was removed by centrifugation at 10,000 *×g* for 60 min at 4 °C. Ni-NTA agarose (2 mL) was added to the supernatant and incubated at 4 °C for 1 h. The resin bound His_6_-tagged proteins were loaded onto a gravity flow column, washed with 0.5 L of wash buffer (50 mM NaH_2_PO_4_, 300 mM NaCl, 20 mM imidazole, 10% glycerol, pH = 8.0), and then eluted with 5 mL of elution buffer (50 mM NaH_2_PO_4_, 300 mM NaCl, 250 mM imidazole, 10% glycerol, pH = 8.0) to desorb the target proteins. Each protein solution was concentrated using a suitable size-exclusion ultrafiltration tube (10 kD for TlnD_TE_, Sfp, and *Sel*Fdx1499; 50 kD for TlnA and its matants, TlnC_ACP*(apo)*-Cy_ and its matants, TlnD_A-PCP*(holo)*_, PbACL891, *Sel*FdR0978, and TlmB’; Amicon, Shanghai, China), and the buffer was exchanged with desalting buffer (50 mM NaH_2_PO_4_, 150 mM NaCl, 10% glycerol, pH = 7.4) using PD-10 desalting column (Beijing, China, GE Healthcare) to remove imidazole. Subsequently, the His_6_-tags and SUMO-tags in the recombinant proteins were removed using thrombin and Ulp1 following the manufacturers’ instructions_79_. Finally, the purified proteins were flash frozen in liquid nitrogen and stored at −80 °C for later use.

### 4.11 Protein concentration determination

The functional concentration of P450 TlnA was determined using the extinction coefficient (ϵ450-490) of 91,000 M^-1^·cm^-1^, which was calculated from the CO-bound reduced difference spectrum, as previously described^80^. The concentrations of other proteins were determined by recording absorbance (A280) using a Nanodrop instrument (Thermo, USA).

### 4.12 Chemical synthesis of compound 7 and D_3_-7

See Supporting Information for the synthetic scheme and NMR characterization details of **7** and D_3_-**7**.

### 4.13 Enzymatic synthesis of 7-CoA and D_3_-7-CoA

To prepare **7**-CoA, a preparative-scale enzymatic reaction (30 mL) in 10 mM NaH_2_PO_4_ buffer (pH = 7.4) containing 2 mM ATP, 2 mM CoASH, 10 mM MgCl_2_, 2 mM DTT, 2 mM **7**, and 50 μM PbACL891 was carried out at 30 °C for 30 min. The reaction was terminated by adding 30 mL of methanol, and the mixture was dried under vacuum. The resulting extract was then subject to HPLC using a YMC-Pack Pro C18 column (10 mm × 250 mm × 5 μM, YMC, Japan). The HPLC program for **7**-CoA purification was as follow: 0 - 1 min, 10 % (ACN in H_2_O with 0.03% formic acid); 2 - 12 min, 10 - 66 %; 12 - 17 min, 100 % and 17 - 23 min, 10 %; 2.5 mL/min, 280 nm. Finally, 8.3 mg of **7**-CoA was obtained after freeze-drying of the collected HPLC fractions. The structure of **7**-CoA was determined by HRESI-MS and NMR analyses (see Supporting Information for details).

To prepare D_3_-**7**-CoA, a preparative-scale enzymatic reaction (15 mL) in 10 mM NaH_2_PO_4_ buffer (pH = 7.4) containing 2 mM ATP, 2 mM CoASH, 10 mM MgCl_2_, 2 mM DTT, 2 mM D_3_-**7**, and 50 μM PbACL891 was carried out at 30 °C for 40 min. The reaction was terminated by adding 30 mL of methanol, and the mixture was dried under vacuum. The resulting extract was then subject to HPLC using a YMC-Pack ODS-A column (10 mm × 250 mm × 5 μM, YMC, Japan). The HPLC program for D_3_-**7**-CoA purification was as follow: 0 - 1 min, 15 % (ACN in H_2_O with 0.03% formic acid as solvent B) in water (as solvent A); 2 - 22 min, 10 - 30 % B; 22 - 27 min, 100 % B and 27 - 32 min, 15 % B; 2.5 mL/min, 280 nm. Finally, 5.4 mg of D_3_-**7**-CoA was obtained after freeze-drying of the collected HPLC fractions. The structure of D_3_-**7**-CoA was determined by HRESI-MS and NMR analyses (see Supporting Information for details).

### 4.14 Chemical synthesis of 6

To prepare compound **6**, 5 mg **7**-CoA (500 μM) was added to 10 mL of (NH_4_)_2_S solution (1 M), and the mixture was stirred at room temperature for 30 min. The reaction mixture was extracted with ethyl acetate three times to gain the crude extract of **6**. The resulting extract was then subject to HPLC using a YMC-Pack Pro C18 column (10 mm × 250 mm × 5 μM, YMC, Japan). The HPLC program for **6** purification was as follows: 0 - 1 min, 40 % (ACN in H_2_O); 2 - 16 min, 40 - 56 %; 17 - 22 min, 100 %, and 22 - 27 min, 40 %; 2.5 mL/min, 230 nm. Finally, 0.7 mg of **6** was obtained after freeze-drying of the collected HPLC fractions. The structure of **6** was identified by HRESI-MS and NMR analyses and compared with the reported spectra (Figures S4, S76, and S77) ^25^.

### 4.15 HPLC-HRESI-MS analysis of ACP/PCP-bound intermediates

Unless otherwise specified, all enzymatic assays for protein mass spectrometric analysis were performed at 30 °C in a 50 μL reaction system consisting of 10 mM NaH_2_PO_4_ (pH = 7.4).

To examine whether _L_-Cys or *S*-methyl-_L_-cysteine could be loaded onto TlnD_A-PCP*(holo)*_, 50 μM purified TlnD_A-PCP*(holo)*_ was incubated with 2 mM _L_-Cys or 2 mM *S*-methyl-_L_-cysteine, 2 mM ATP, 10 mM MgCl_2_, and 2 mM DTT for 2 h.

To detect the online catalytic intermediates of TlnC_ACP-Cy_ mutants, 50 μM TlnD_A-PCP*(holo)*_ was incubated with 2 mM _L_-Cys, 500 μM **7**-CoA, and 10 μM TlnC_ACP*(apo)*-Cy_ wild type or its mutants (TlnC_ACP*(apo)*-Cy-D140A_, TlnC_ACP*(apo)*-Cy-N352A_, TlnC_ACP*(apo)*-Cy-N352T,_ or TlnC_ACP*(apo)*-Cy-D394A_) in the presence of 2 mM DTT, 2 mM ATP, and 10 mM MgCl_2_ for 15 min.

To examine whether **7** could be loaded onto TlnC_ACP*(apo)*-Cy_, 50 μM TlnC_ACP*(apo)*-Cy_ was reacted with 5 μM Sfp, 500 μM **7**-CoA in the presence of 10 mM MgCl_2_ and 2 mM DTT for 15 min.

To examine whether TlnA could catalyze an online reaction, 2 μM TlnA, 5 μM *sel*Fdx1499, 10 μM *sel*FdR0978 were reacted with 50 μM TlnC_ACP*(apo)*-Cy_, 5 μM Sfp, 500 μM **7**-CoA in the presence of 10 mM MgCl_2_, 2 mM DTT, and 2 mM NADPH, and the reaction mixture was incubated for 30 min.

Afterwards, the reactant mixtures were centrifuged (12,000 *×g*) at 4 °C for 10 min, and the supernatants were used for HPLC-ESI-MS analysis. The HPLC program was as follow: 0 - 1 min, 5 % (ACN in H_2_O with 0.1% formic acid); 2 - 15 min, 5 - 90 %; 15 - 18 min, 90 % and 19 - 25 min, 5%; 1 mL/min, 280 nm, 70 °C.

### 4.16 *In vitro* enzymatic assays

Unless otherwise specified, all enzymatic assays in this part were performed at 30 °C in a 100 μL reaction system consisting of 10 mM NaH_2_PO_4_ (pH = 7.4), and the boiling-inactivated enzymes were used for negative control reactions.

The one-pot reaction for **1** biosynthesis contained 10 μM TlnC_ACP*(apo)*-Cy_, 10 μM TlnD_A-PCP*(holo)*_, 2 μM TlnD_TE_ (*optional*), 2 μM TlnA, 5 μM *sel*Fdx1499, 10 μM *sel*FdR0978, 2 mM ATP, 10 mM MgCl_2_, 500 μM **7**-CoA, 2 mM DTT, 2 mM _L_-Cys, and 2 mM NADPH. The reaction was carried out for 4 h.

To examine whether **6** could be converted by TlnA, 500 μM **6** was incubated with 2 μM TlnA, 5 μM *sel*Fdx1499, 10 μM *sel*FdR0978, and 2 mM NADPH for 2 h.

To examine whether the *in situ* produced **9** could be converted by TlnA, 2 μM TlnA, 5 μM *sel*Fdx1499, 10 μM *sel*FdR0978 were co-incubated with 500 μM **7**-CoA in the presence of 20 mM (NH_4_)_2_S and 2 mM NADPH for 2 h.

To analyze the activity of TlnA and its mutants with compound **7** as substrate, 2 μM TlnA or its mutants, 5 μM *sel*Fdx1499, 10 μM *sel*FdR0978, and 2 mM NADPH were co-incubated with 200 μM **7** for 2 h. The enzyme reaction of TlnA and its mutants with *in situ* produced compound **9** contained 10 μM TlnC_ACP*(apo)*-Cy_, 10 μM TlnD_A-PCP*(holo)*_, 2 μM TlnA or its mutants, 5 μM *sel*Fdx1499, 10 μM *sel*FdR0978, 2 mM ATP, 10 mM MgCl_2_, 500 μM **7**-CoA, 2 mM _L_-Cys, and 2 mM NADPH for 2 h.

To reconstitute the sulfur-incorporation reaction co-mediated by TlnC_ACP-Cy_ and TlnD_A-PCP_, 10 μM TlnC_ACP*(apo)*-Cy_, 10 μM TlnD_A-PCP*(holo)*_, 500 μM **7**-CoA, and 2 mM _L_-Cys were incubated with 2 mM ATP, 10 mM MgCl_2_, and 2 mM DTT for 2 h. To analyze the activity of TlnC_ACP-Cy_ mutants, the wild-type TlnC_ACP*(apo)*-Cy_ was individually replaced by its mutants (TlnC_ACP*(apo)*-Cy-D140A_, TlnC_ACP*(apo)*-Cy-N352A_, TlnC_ACP*(apo)*-Cy-N352T,_ or TlnC_ACP*(apo)*-Cy-D394A_) in the above reaction system.-In the cysteamine derivatization reaction, 10 mM cysteamine was further added to the reaction.

Afterwards, each reaction described above was terminated by adding 200 μL methanol, and the mixture was then highspeed centrifuged (12,000 *×g*, 10 min, 16 °C) to remove the metamorphic proteins. Finally, the supernatants were analyzed with HPLC or HPLC-HRMS. The HPLC program was follow: 0 - 1 min, 10 % (ACN in H_2_O with 10 mM ammonium acetate for detection of **7**-CoA); 2 - 25 min, 10 - 80 %; 26 - 30 min, 100 % and 30 - 35 min, 10%; 1 mL/min, 280 nm.

### 4.17 Preparation of the decarboxylation product 11

To prepare **11**, a preparative-scale enzymatic reaction (15 mL) in 10 mM NaH_2_PO_4_ buffer (pH = 7.4) containing 2 μM TlnA, 5 μM *sel*Fdx1499, 10 μM *sel*FdR0978, 2 mM NADPH, and 2 mM compound **7** was carried out at 30°C for 4 h. The reaction was terminated by adding 15 mL of methanol, and the mixture was dried under vacuum. The resulting extract was then subject to HPLC using a YMC-Pack Pro C18 column (10 mm × 250 mm × 5 μM, YMC, Japan). The HPLC program for **11** purification was as follows: 0 - 1 min, 34 % (ACN in H_2_O); 2 - 20 min, 34 - 80 %; 20 - 25 min, 100 % and 25 - 30 min, 34 %; 2.5 mL/min, 280 nm. Finally, 1.4 mg of purified **11** was obtained after freeze-drying of the collected HPLC fractions. The structure of **11** was determined by HRESI-MS and NMR analyses (see Supporting Information for details).

### 4.18 Quantitative mass spectrometry analysis of the formation of 1 and D_2_-1 in TlnA-(NH_4_)_2_S mediated reaction

Quantitative mass spectrometry analysis was performed on an Ultra-High Performance liquid chromatography system (SCIEX, ExionLC, UHPLC) coupled with a triple quadrupole mass spectrometer (SCIEX Triple Quad 5500+ QTrap Ready). Chromatographic separation of **1** and D_2_-**1** was achieved on a Kinetex C18 100 Å column (100 mm ×2.1 mm, 2.6 µm). The two eluents were 0.1% formic acid in water (A) and 0.1% formic acid in acetonitrile (B). The mobile phase was delivered at a flow rate of 0.3 mL/min using a gradient of A and B as follows: 0-5 min, 5-98 % B; 5-7 min, 98 % B; 7-10 min, 5 % B. The mass spectrometer was operated in positive electrospray ionization (ESI) using the multiple reaction (MRM) mode with a dwell time of 20 ms per transition. The source/gas-dependent parameters were optimized and set as follows: curtain gas 35 psi; collision gas 9 psi; spray voltage 4500 V; temperature 550 °C; ion source gas 1 (GS1) 60 psi; ion source gas 2 (GS2) 60 psi. The mass transitions and compound-dependent parameters were optimized using the authentic standards and are listed in Table S3.

### 4.19 Isotope labeling experiments of the TlnA-mediated hydroxylation reaction

To understand the catalytic mechanism of TlnA, ^18^O-labeled water (≥98% labeled, Tenglong Weibo, Qingdao) and ^18^O_2_ (≥98% labeled, Delin, Shanghai) were used to replace normal water and O_2_ in the reaction system, respectively.

For ^18^O_2_ labeling experiments, the reaction was performed at 30 °C in 200 μL reaction buffer (H_2_O) in an Ampoule bottle (2 mL) containing 2 μM TlnA, 5 μM *sel*Fdx1499, 10 μM *sel*FdR0978, and 500 μM **7**. Then the bottle was purged with nitrogen and sealed with a bottle cap. ^18^O_2_ was introduced into the bottle from the compressed gas bag using a syringe needle. Finally, 2 mM NADPH was added to the reaction mixture using a microsyringe to start the reaction. After incubation at 30 °C for 2 h, 200 μL of methanol was added using a microsyringe to terminate the reactions.

For H_2_^18^O labeling experiments, the reaction was performed at 30 °C in 200 μL reaction buffer (H_2_^18^O) in an Ampoule bottle (2 mL) containing 2 μM TlnA, 5 μM *sel*Fdx1499, 10 μM *sel*FdR0978, and 500 μM **7**. After incubation for 2 h, 200 μL of methanol was added to quench the reaction. HRESI-LCMS analysis was carried out to detect the products using the abovementioned method.

### 4.20 Protein structure prediction and molecular docking

The 3D structures of TlnA, TlnC_Cy_ and TlnD_PCP_, TlnC_ACP-Cy_-TlnA and TlnD-TlnA monomers were predicted using AlphaFold3 program,^49^ and the structures were optimized using an Amber force field with default parameters^81^. With the predicted monomer structures, the structural model (Per-residue Local Distance Difference Test (pLDDT) = 90.536) was selected based on the pLDDT score for further analysis. The compound **14** was manually connected to S138 of TlnD_PCP_ monomer by AutoDock Vina to build the model of TlnD_PCP_ tethered **14**. Subsequently, substrate **9** was docked into TlnA, and TlnD_PCP_ tethered **14** was docked into TlnC_Cy_. The AutoDock Vina^50^ molecular docking algorithm was used to establish the binding mode between small molecules and proteins. The generated binding modes were evaluated using the AI scoring algorithm of OnionNet-SFCT^82^. During this process, a pocket size of 20 Å was set with the Fe atom of heme as the center, and the exhaustness parameter was kept at 64, while other parameters were maintained at default values. For the OnionNet-SFCT scoring algorithm, weightings of 0.2 (for Vina score) and 0.8 (for correction terms) were assigned.

### 4.21 QM/MM calculations for enzymatic reactions

All QM/MM calculations were performed using ChemShell,^83, 84^ combining Turbomole^85^ for the QM region and DL_POLY^86, 87^ for the MM region. The polarizing effect of the protein environment on the QM region was treated with the electrostatic embedding scheme^88^. During the QM/MM calculations, the hybrid B3LYP^89^ functional was used for the QM region, while the same Amber force field was used for the MM region. The double-*ζ* basis set def2-SVP^90^(B1) was used for geometry optimizations, while the def2-TZVP(B2) basis set was employed for the single-point energy calculations. The dispersion corrections computed with Grimme’s D3^91, 92^ method were considered in all QM/MM calculations. For whole reactions, the QM region consisted of the substrate and HEM (see Supporting Information for details).

### 4.22 The derivatization assay

All one-pot reactions for derivatization assay were performed in a 100 μL reaction system consisting of 10 mM NaH_2_PO_4_ (pH = 7.4) containing 10 μM TlnC_ACP*(apo)*-Cy_, 10 μM TlnD_A-PCP*(holo)*_, 200 μM **7**-CoA, 2 mM _L_-Cys, 500 μM ATP, 10 mM MgCl_2_, and 2 mM DTT at 30 °C for 4 h.

For 4-FPH derivatization, 100 µL 4-FPH solution (0.1 mg/mL in 50% ethanol) was added to the one-pot reaction described above. After vortex-oscillation for 30 s, the reaction mixture was further incubated at 50 °C for 3 h. After high-speed centrifugation (12,000 *×g*, 10 min, 16 °C), the supernatant was analyzed by HPLC-HRESI-MS. The HPLC program was as follows: 0 - 1 min, 10 % (ACN in H_2_O with 0.1% formic acid); 2 - 25 min, 10 - 80 %; 25 - 30 min, 100 % and 30 - 35 min, 10%; 1 mL/min, 280 nm.

For OPD derivatization, 100 μL of 12 mM OPD in 3 N HCl was added to the one-pot reaction described above. After vortex-oscillation for 30 s, the reaction mixture was further incubated at 100 °C for 30 min. After high-speed centrifugation (12,000 *×g*, 10 min, 16 °C), the supernatant was analyzed by HPLC. The HPLC program was as follow: 0 - 1 min, 10 % (ACN in H_2_O); 2 - 25 min, 10 - 80 %; 26 - 30 min, 100 % and 30 - 35 min, 10%; 1 mL/min, 280 nm.

For DC derivatization, 100 µL DC solution (5 mg/mL in ACN) and 200 µL carbonate/sodium bicarbonate buffer (pH = 11.5) were added to the one-pot reaction described above. After vortex-oscillation for 30 s, the reaction mixture was incubated at 60 °C under lightless conditions for 20 min. The dansyl chloride-alanine derivative was extracted by adding 400 μL of ethyl acetate. After high-speed centrifugation (12,000 *×g*, 10 min, 16 °C), the supernatant was dried and dissolved in 100 μL methanol for HPLC-HRESI-MS analysis. The HPLC program for detecting alanine derivative was as follow: 0 - 1 min, 60 % (ACN in H_2_O with 0.1% formic acid); 2 - 30 min, 60 - 100 %; 30 – 35 min, 100 % and 35 - 40 min, 60%; 1 mL/min, 280 nm.

For HP derivatization, 10 µL riphenylphosphine (10 mM in ACN), 10 µL 2,2’-dipyridyl disulfide (10 mM in ACN), and 10 µL HP (10 mg/mL) were added to the one-pot reaction described above. After vortex-oscillation for 30 s, the reaction mixture was incubated at 60 °C for 10 min. After high-speed centrifugation (12,000 *×g*, 10 min, 16 °C), the supernatant was analyzed by HPLC-HRESI-MS analysis. The HPLC program for detecting alanine derivative was as follow: 0 - 1 min, 2 % (ACN in H_2_O with 0.1% formic acid); 2 - 30 min, 2 - 60 %; 30 - 35 min, 100 % and 35 - 40 min, 2%; 1 mL/min, 280 nm.

### 4.23 Protein conformation analysis using circular dichroism spectrum (CD)

Protein samples were diluted to a concentration of 10 μM, and the CD spectra were collected using a wavelength range of 190-260 nm with an interval of 0.5 nm, at 25 °C in a 200 μL cuvette. The desalting buffer was used to dissolve proteins and served as a background to be subtracted from each sample spectrum.

### 4.24 ^13^C NMR-tracer experiments for the identification of the C_3_ product

For the ^13^C NMR time-course experiments, the 1.5 mL reaction system contained 500 μM _L_-[1,2,3-^13^C3,^15^N] cysteine, 800 μM **7**-CoA, 20 μM TlmB’, ATP, MgCl_2_, and DTT in either 50 mM NaH_2_PO_4_ or 50 mM Tris-HCl buffer. The enzymatic reactions were divided into three 0.5 mL aliquots and quenched with an equal volume of methanol at three time points of 1, 30, and 240 min. To increase the C_3_ product concentration while keeping the salt concentration below 200 mM (essential for NMR detection sensitivity), all samples were concentrated 4-fold through lyophilization and re-dissolution in D_2_O prior to ^13^C-NMR analysis.

### 4.25 Isothermal titration calorimetry (ITC)

The ITC data were recorded on a MicroCal PEAQ-ITC (Malvern Panalytical) equipment at 25°C. Protein samples (TlnA and TlnC_ACP*(apo)*-Cy_) were exchanged to an ITC buffer (containing 50 mM NaH_2_PO_4_, 150 mM NaCl, and 5% Glycerol, pH = 7.4) and centrifuged at 12,000 *×g* for 10 min, before being subjected to the measurements. For titration, after an initial delay of 60 s, 0.4 μL of TlnA (220 μM) was first injected into the cell (containing 20 μM TlnC_ACP-Cy_), and then another 18 times of 2.0 μL injections were followed with a spacing time of 150 s to allow for equilibration. The cell was stirred at 750 rpm, and the reference power was 10 ucal/s. TlnA was injected into the ITC buffer as a blank titration.

### 4.26 Protein-protein interaction analysis by size-exclusion chromatography (SEC)

The high-purity TlnA was incubated with SUMO-TlnC_ACP-Cy_ at 30°C for 0.5 h for full interactions. After that, the single protein TlnA and SUMO-tagged TlnC_ACP-Cy_, as well as their mixture, were analyzed by SEC using a SuperdexTM 75 Increase 10/300 GL in the buffer containing 50 mM NaH_2_PO_4_, 150 mM NaCl, and 10% Glycerol with a flow rate of 0.4 mL/min.

### 4.27 Protein-protein interaction analysis by native-PAGE

TlnA, SUMO-tagged TlnC_ACP-Cy_, and their 1:1 mixture were analyzed by Native-PAGE Plastic gel (Tris-Gly 4-20%, WSHTbio).

### 4.28 NMR data of 1-4, 6, 7-CoA, D_3_-7-CoA, and 11

**1**: ^1^H NMR (600 MHz, CD_3_OD) δ 6.37 (dd, *J* = 17.4, 10.7 Hz, 1H), 5.63 (s, 1H), 5.27 (d, *J* = 17.4 Hz, 1H), 5.05 (d, *J* = 10.7 Hz, 1H), 1.81 (s, 3H), 1.71 (s, 3H), 1.70 (s, 3H). ^13^C NMR (151 MHz, CD_3_OD) δ 197.55, 182.88, 142.22, 140.50, 131.43, 113.74, 110.25, 56.39, 49.57, 49.43, 49.28, 49.14, 49.00, 48.86, 48.72, 48.57, 30.20, 11.96, 7.65.

**2**: ^1^H NMR (600 MHz, CD_3_OD) δ 6.38 (dd, *J* = 17.3, 10.7 Hz, 1H), 5.62 (s, 1H), 5.25 (d, *J* = 17.4 Hz, 1H), 5.04 (d, *J* = 10.7 Hz, 1H), 2.14 – 2.01 (m, 2H), 1.71 (s, 3H), 1.69 (s, 3H), 0.90 (t, *J* = 7.1 Hz, 3H).

**3**: ^1^H NMR (600 MHz, CD_3_OD) δ 6.40 (dd, *J* = 17.4, 10.7 Hz, 1H), 5.63 (s, 1H), 5.30 (d, *J* = 17.4 Hz, 1H), 5.08 (d, *J* = 10.7 Hz, 1H), 2.24 (dd, *J* = 12.5, 7.2 Hz, 2H), 1.83 (s, 3H), 1.75 (s, 3H), 1.04 (t, *J* = 7.5 Hz, 3H).

**4**: ^1^H NMR (600 MHz, CD_3_OD) δ 6.41 (dd, *J* = 17.4, 10.7 Hz, 1H), 5.63 (s, 1H), 5.29 (d, *J* = 17.4 Hz, 1H), 5.08 (d, *J* = 10.7 Hz, 1H), 2.28 – 2.21 (m, 2H), 2.10 (tq, *J* = 14.0, 7.0 Hz, 2H), 1.03 (t, *J* = 7.5 Hz, 3H), 0.94 (t, *J* = 7.2 Hz, 3H).

**6**: ^1^H NMR (600 MHz, CDCl_3_) δ 5.45 (d, *J* = 6.6 Hz, 1H), 4.53 (d, *J* = 4.0 Hz, 1H), 3.91 (d, *J* = 6.6 Hz, 1H), 3.03 (dd, *J* = 7.1, 4.1 Hz, 1H), 1.68 (s, 2H), 1.68 – 1.66 (m, 5H), 1.32 (d, *J* = 6.5 Hz, 3H), 1.13 (d, *J* = 7.1 Hz, 3H). ^13^C NMR (151 MHz, CDCl_3_) δ 205.50, 196.68, 129.93, 124.88, 60.48, 51.06, 45.96, 15.25, 13.62, 11.81, 8.49.

**7**-CoA: ^1^H NMR (600 MHz, D_2_O) δ 8.53 (s, 1H), 8.30 (d, *J* = 1.8 Hz, 1H), 7.08 (d, *J* = 3.1 Hz, 1H), 6.07 (s, 1H), 5.83 (p, *J* = 6.6 Hz, 1H), 4.75 (dd, *J* = 9.5, 5.5 Hz, 1H), 4.68 – 4.64 (m, 1H), 4.47 (s, 1H), 4.14 (s, 2H), 3.89 (s, 1H), 3.74 (dd, *J* = 9.4, 4.0 Hz, 1H), 3.51 – 3.46 (m, 1H), 3.29 (d, *J* = 2.7 Hz, 2H), 3.23 – 3.16 (m, 2H), 2.89 (d, *J* = 5.9 Hz, 2H), 2.26 (t, *J* = 6.6 Hz, 2H), 1.76 (s, 3H), 1.74 (s, 3H), 1.63 (d, *J* = 6.8 Hz, 3H), 1.19 (d, *J* = 6.8 Hz, 3H), 0.80 (s, 3H), 0.68 (s, 3H). ^13^C NMR (151 MHz, D_2_O) δ 201.2, 200.8, 174.3, 173.8, 149.5, 148.6, 148.5, 144.7, 142.7, 136.4, 133.5, 133.4, 118.5, 87.4, 83.9, 74.5, 73.8, 72.1, 65.6, 55.1, 38.5, 35.6, 35.3, 28.3, 21.4, 18.2, 15.1, 14.7, 14.0, 12.8.

D_3_-**7**-CoA: ^1^H NMR (600 MHz, D_2_O) δ 8.44 (s, 1H), 8.14 (d, *J* = 3.4 Hz, 1H), 7.06 (d, *J* = 7.5 Hz, 1H), 6.06 (d, *J* = 5.1 Hz, 1H), 5.81 (d, *J* = 9.9 Hz, 1H), 4.50 (s, 1H), 4.17 (s, 2H), 3.93 (s, 1H), 3.75 (d, *J* = 6.3 Hz, 1H), 3.48 (d, *J* = 6.4 Hz, 1H), 3.33 (dd, *J* = 12.2, 6.2 Hz, 2H), 3.22 (dd, *J* = 12.6, 7.1 Hz, 2H), 3.02 – 2.86 (m, 2H), 2.30 (t, *J* = 6.5 Hz, 2H), 1.79 (s, 3H), 1.75 (d, *J* = 5.2 Hz, 3H), 1.24 (dd, *J* = 6.7, 3.8 Hz, 3H), 0.81 (d, *J* = 9.4 Hz, 3H), 0.67 (d, *J* = 3.6 Hz, 3H). ^13^C NMR (151 MHz, D_2_O) δ 201.1, 200.7, 174.6, 173.8, 155.4, 152.7, 149.2, 148.1, 139.7, 136.5, 133.7, 132.2, 119.9, 86.46, 83.6, 74.0, 73.8, 71.8, 65.4, 54.6, 38.4, 35.3, 35.3, 28.2, 20.8, 18.2, 15.0, 14.5, 14.4, 12.5.

**11**: ^1^H NMR (600 MHz, DMSO-*d*_*6*_) δ 7.02 (s, 1H), 5.75 (t, *J* = 6.2 Hz, 1H), 4.10 (t, *J* = 5.6 Hz, 2H), 2.72 (q, *J* = 7.2 Hz, 2H), 1.86 (d, *J* = 1.1 Hz, 3H), 1.81 (d, *J* = 0.8 Hz, 3H), 0.96 (t, *J* = 7.2 Hz, 3H). ^13^C NMR (151 MHz, DMSO-*d*_*6*_) δ 202.63, 142.05, 137.04, 134.17, 132.17, 58.03, 30.17, 16.67, 13.08, 8.99.

## ASSOCIATED CONTENT

### Supporting Information

The Supporting Information contains additional methods, tables, schemes, and figures. HR-MS data and NMR spectra are available free of charge on the ACS Publications website at DOI:XXX (PDF)

## AUTHOR INFORMATION

Authors

**Jiawei Guo** - *State Key Laboratory of Microbial Technology, Shandong University, Qingdao, Shandong, 266237, China*

**Qiaoyu Zhang** - *State Key Laboratory of Physical Chemistry of Solid Surfaces and Fujian Provincial Key Laboratory of Theoretical and Computational Chemistry, College of Chemistry and Chemical Engineering, Xiamen University, Xiamen, Fujian, 361005, China*

**Yang Shen** - *State Key Laboratory of Physical Chemistry of Solid Surfaces, Key Laboratory of Chemical Biology of Fujian Province, iCHEM, College of Chemistry and Chemical Engineering, Xiamen University, Xiamen, Fujian 361005, China*

**Fangyuan Cheng** - *State Key Laboratory of Microbial Technology, Shandong University, Qingdao, Shandong, 266237, China*; *Laboratory for Marine Biology and Biotechnology, Qingdao National Laboratory for Marine Science and Technology, Qingdao, Shandong, 266237, China*

**Moli Sang** - *State Key Laboratory of Microbial Technology, Shandong University, Qingdao, Shandong, 266237, China*

**Xuan Wang** - *State Key Laboratory of Microbial Technology, Shandong University, Qingdao, Shandong, 266237, China*

**Yunjun Pan** - *State Key Laboratory of Microbial Technology, Shandong University, Qingdao, Shandong, 266237, China*

**Mingyu Liu -** *State Key Laboratory of Microbial Technology, Shandong University, Qingdao, Shandong, 266237, China*

**Hao-Bing Yu** - *Department of Marine Biomedicine and Polar Medicine, Naval Medical Center, Naval Medical University, Shanghai, 200433, China*

**Bo Hu** - *Department of Marine Biomedicine and Polar Medicine, Naval Medical Center, Naval Medical University, Shanghai, 200433, China*

**Sheng Wang** - *Shanghai Zelixir Biotech Company Ltd*., *Shanghai 200030, China*

**Liangzhen Zheng** - *Shanghai Zelixir Biotech Company Ltd*., *Shanghai 200030, China*

**Ce Geng** - *Shandong Provincial Key Laboratory of Synthetic Biology, Qingdao Institute of Bioenergy and Bioprocess Technology, Chinese Academy of Sciences, Qingdao, Shandong, 266101, China*

**Chaofan Yang** - *State Key Laboratory of Microbial Technology, Shandong University, Qingdao, Shandong, 266237, China*

**Lianzhong Luo** - *Xiamen Key Laboratory of Marine Natural Product Resources, Xiamen Medical College, Xiamen, 361023, China*

**Gang Zhang** - *Xiamen Key Laboratory of Marine Natural Product Resources, Xiamen Medical College, Xiamen, 361023, China*

**Lei Du** - *State Key Laboratory of Microbial Technology, Shandong University, Qingdao, Shandong, 266237, China*

**Yuanning Li** - *State Key Laboratory of Microbial Technology, Shandong University, Qingdao, Shandong, 266237, China*

**Wei Zhang** - *State Key Laboratory of Microbial Technology, Shandong University, Qingdao, Shandong, 266237, China*

## Author Contributions

## Notes

The authors declare no competing financial interest.

## ACKNOWLEDGMENTS

The authors thank Professor Song Xue from Dalian University of Technology for her gift of the *Streptomyces* sp. S6043a strain. The authors are grateful to Professor Hung-wen Liu from University of Texas at Austin and Professor Wen Liu from Shanghai institute of Organic Chemistry, Chinese Academy of Sciences, for their helpful discussions. The authors also thank Jingyao Qu, Guannan Lin, Jing Zhu, Zhifeng Li, Haiyan Sui and Xiangmei Ren from the State Key Laboratory of Microbial Technology at Shandong University for their guidance and assistance in HPLC, HPLC-MS, ITC and NMR analysis.

This work was supported by the National Key R&D Program of China (2023YFA0915500), the National Natural Science Foundation of China (32470043, 323B2049, 32025001, 22237004, 22471227, 22301251), the Taishan Scholars Program (tsqn202408318), the Shandong Provincial Natural Science Foundation (ZR2023ZD50), the China National Postdoctoral Program for Innovative Talents (BX20240210), the State Key Laboratory of Microbial Technology Open Projects Fund (M2023-01), and Fujian Province Universities and Colleges Technology and Engineering Center for Marine Biomedical Resource, Xiamen Medical College (xmmc-mnpr-2022001).

## Notes

### Competing Interest Statement

The authors have declared no competing interest.

### Summary of Updates

Authors and affiliations updated; Figures 1-3 updated; Sections on 2.4 and 2.5 updated to clarify the mechanism of TlnA and TlnCCy.

## REFERENCES

(1) Cao, X.; Cao, L.; Zhang, W.; Lu, R.; Bian, J. S.; Nie, X. Therapeutic Potential of Sulfur-Containing Natural Products in Inflammatory Diseases. Pharmacol. Ther. 2020, 216, 107687–107700.

(2) Francioso, A.; Baseggio Conrado, A.; Mosca, L.; Fontana, M. Sulfur-Containing Marine Natural Products as Leads for Drug Discovery and Development. Curr. Opin. Chem. Biol. 2023, 75, 102330102345.

(3) Liu, M.; Zhang, X.; Li, G. Structural and Biological Insights into the Hot-Spot Marine Natural Products Reported from 2012 to 2021. Chin. J. Chem. 2022, 40 (15), 1867–1889.

(4) Chatterjee, A; Cao, L.; Zhang, W.; Lu, R.; Bian, J. S.; Nie, X. Saccharomyces Cerevisiae THI4p is a Suicide Thiamine Thiazole Synthase. Nature 2011, 478, 542–546.

(5) Meng, S.; Steele, A. D.; Yan, W.; Pan, G.; Kalkreuter, E.; Liu, Y. C.; Xu, Z.; Shen, B. Thiocysteine Lyases as Polyketide Synthase Domains Installing Hydropersulfide into Natural Products and A Hydropersulfide Methyltransferase. Nat. Commun. 2021, 12, 5672–5682.

(6) Cao, M.; Zheng, C.; Yang, D.; Kalkreuter, E.; Adhikari, A.; Liu, Y. C.; Rateb, M. E.; Shen, B. Cryptic Sulfur Incorporation in Thioangucycline Biosynthesis. Angew. Chem. Int. Ed. Engl. 2021, 60, 7140–7147.

(7) Zhao, Q.; Wang, M.; Xu, D.; Zhang, Q.; Liu, W. Metabolic Coupling of Two Small-Molecule Thiols Programs the Biosynthesis of Lincomycin A. Nature 2015, 518, 115–119.

(8) Sasaki, E.; Zhang, X.; Sun, H. G.; Lu, M. Y. Liu, T. L.; Ou, A.; Li, J. Y.; Chen, Y. H.; Ealick, S. E.; Liu, H. W. Co-Opting Sulphur-Carrier Proteins from Primary Metabolic Pathways for 2-Thiosugar Biosynthesis. Nature 2014, 510, 427–431.

(9) Zhang, X.; Xu, X.; You, C.; Yang, C.; Guo, J.; Sang, M.; Geng, C.; Cheng, F.; Du, L.; Shen, Y.; Wang, S.; Lan, H.; Yang, F.; Li, Y.; Tang, Y.-J.; Zhang, Y.; Bian, X.; Li, S.; Zhang, W. Biosynthesis of Chuangxinmycin Featuring a Deubiquitinase-Like Sulfurtransferase. Angew. Chem. Int. Ed. Engl. 2021, 60, 24418–24423.

(10) Steele, A. D.; Kiefer, A. F.; Shen, B. The Many Facets of Sulfur Incorporation in Natural Product Biosynthesis. Curr. Opin. Chem. Biol. 2023, 76, 102366–102374.

(11) Oishi, H.; Noto, T.; Sasaki H.; Suzuki, K.; Hayashi, T.; Okazaki, H.; Ando, K.; Sawada, M. Thiolactomycin, a new antibiotic I. Taxonomy of the Producing Organism, Fermentation and Biological Properties. J. Antibiot. 1982, 35, 391–395.

(12) Sasaki, H.; Noto, T.; Sasaki, H.; Suzuki, K.; Hayashi, T.; Okazaki, H.; Ando, K.; Sawada, M. Thiolactomycin, a New Antibiotic II. Structure Elucidation. J. Antibiot. 1982, 35, 396–400.

(13) Price, A. C.; Choi, K. H.; Heath, R. J.; Li, Z.; White, S. W.; Rock, C. O. Inhibition of Beta-Ketoacyl-Acyl Carrier Protein Synthases by Thiolactomycin and Cerulenin. J. Biol. Chem. 2001, 276, 6551–6559.

(14) Miyakawa, S.; Suzuki, K.; Noto, T.; Harada, Y.; Okazaki, H. Thiolactomycin, a New Antibiotic IV. Biological Properties and Chemotherapeutic Activity in Mice. J. Antibiot. 1982, 35, 411–419.

(15) Noto, T.; Miyakawa, S.; Oishi, H.; Endo, H.; Okazaki, H. Thiolactomycin, a New Antibiotic III. in Vitro Antibacterial Activity. J. Antibiot. 1982, 35, 401–410.

(16) Machutta, C. A.; Bommineni, G. R.; Luckner, S. R.; Kapilashrami, K.; Ruzsicska, B.; Simmerling, C.; Kisker, C.; Tonge, P. Slow Onset Inhibition of Bacterial ?-Ketoacyl-Acyl Carrier Protein Synthases by Thiolactomycin. J. Biol. Chem. 2010, 285, 6161–6169.

(17) Dormann, K. L.; Brückner, R. Variable Synthesis of the Optically Active Thiotetronic Acid Antibiotics Thiolactomycin, Thiotetromycin, and 834-B1. Angew. Chem. Int. Ed. 2007, 46, 11601163.

(18) Ohata, K.; Terashima, S. Efficient Synthesis and Biological Activity of Enantiomeric Pairs of Thiolactomycin and Its 3-Demethyl Derivative. Tetrahedron 2009, 65, 2244–2253.

(19) Brown, M. S.; Akopiants, K.; Resceck, D. M.; McArthur, H. A.; McCormick, E.; Reynolds, K. A. Biosynthetic Origins of the Natural Product, Thiolactomycin: a Unique and Selective Inhibitor of Type II Dissociated Fatty Acid Synthases. J. Am. Chem. Soc. 2003, 125, 10166–10167.

(20) Tang, X.; Li, J.; Millan-Aguinaga, N.; Zhang, J. J.; O’Neill, E. C.; Ugalde, J. A.; Jensen, P. R.; Mantovani, S. M.; Moore, B. S. Identification of Thiotetronic Acid Antibiotic Biosynthetic Pathways by Target-Directed Genome Mining. ACS Chem. Biol. 2015, 10, 2841–2849.

(21) Tao, W.; Yurkovich, M. E.; Wen, S.; Lebe, K. E.; Samborskyy, M.; Liu, Y.; Yang, A.; Liu, Y.; Ju, Y.; Deng, Z.; Tosin, M.; Sun, Y.; Leadlay, P. F. A Genomics-Led Approach to Deciphering the Mechanism of Thiotetronate Antibiotic Biosynthesis. Chem. Sci. 2016, 7, 376–385.

(22) Havemann, J.; Yurkovich, M. E.; Jenkins, R.; Harringer, S.; Tao, W.; Wen, S.; Sun, Y.; Leadlay, P. F.; Tosin, M. Chemical Probing of Thiotetronate Bio-assembly. Chem. Commun. 2017, 53, 1912–1915.

(23) Yurkovich, M. E.; Jenkins, R.; Sun, Y.; Tosin, M.; Leadlay, P. F. The Polyketide Backbone of Thiolactomycin is Assembled by an Unusual Iterative Polyketide Synthase. Chem. Commun. 2017, 53, 2182–2185.

(24) Li, J.; Tang, X.; Awakawa, T.; Moore, B. S. Enzymatic C-H Ox-idation-Amidation Cascade in the Production of Natural and Unnatural Thiotetronate Antibiotics with Potentiated Bioactivity. Angew. Chem. Int. Ed. Engl. 2017, 56, 12234–12239.

(25) Tang, X.; Li, J.; Moore, B. S. Minimization of the Thiolactomycin Biosynthetic Pathway Reveals that the Cytochrome P450 Enzyme TlmF is Required For Five-Membered Thiolactone Ring Formation. Chembiochem 2017, 18, 1072–1076.

(26) Zhang, X. W.; Cheng, F. Y.; Guo, J. W.; Zheng, S. M.; Wang, X.; Li, S. Y., Enzymatic synthesis of organoselenium compounds via C-Se bond formation mediated by sulfur carrier proteins. Nat. Synth. 2024, 3, 477–487.

(27) Xin, Y. J.; Kanagasabhapathy, M.; Janussen, D.; Xue, S.; Zhang, W. Phylogenetic Diversity of Gram-Positive Bacteria Cultured from Antarctic Deep-Sea Sponges. Polar. Biol. 2011, 34, 1501–1512.

(28) Blin, K.; Shaw, S.; Kloosterman, A. M.; Charlop-Powers, Z.; Van Wezel, G. P.; Medema, M. H.; Weber, T. antiSMASH 6.0: Improving Cluster Detection and Comparison Capabilities. Nucleic Acids Res. 2021, 49, W29–W35.

(29) Gnann, A. D.; Marincin, K.; Frueh, D. P.; Dowling, D. P. Dynamics and Mechanistic Interpretations of Nonribosomal Peptide Synthetase Cyclization Domains. Curr. Opin. Chem. Biol. 2023, 72, 102228.

(30) Fortinez, C. M.; Bloudoff, K.; Harrigan, C.; Sharon, I.; Strauss, M.; Schmeing, T. M. Structures and Function of a Tailoring Oxidase in Complex With a Nonribosomal Peptide Synthetase Module. Nat. Commun. 2022, 13, 548–560.

(31) Katsuyama, Y.; Sone, K.; Harada, A.; Kawai, S.; Urano, N.; Adachi, N.; Moriya, T.; Kawasaki, M.; Shin-Ya, K.; Senda, T.; Ohnishi, Y. Structural and Functional Analyses of the Tridomain-Nonribosomal Peptide Synthetase Fmoa3 for 4-Methyloxazoline Ring Formation. Angew. Chem. Int. Ed. 2021, 60, 7694–7699.

(32) Bloudoff, K.; Schmeing, T. M. Structural and Functional Aspects of the Nonribosomal Peptide Synthetase Condensation Domain Superfamily: Discovery, Dissection and Diversity. Biochim. Biophys. Acta (BBA) - Proteins Proteomics 2017, 865, 1587–1604.

(33) Huang, H.; Zheng, G.; Jiang, W.; Hu, H.; Lu, Y. One-Step High-Efficiency CRISPR/Cas9-Mediated Genome Editing in Streptomyces. Acta Biochim. Biophys. Sin. 2015, 47, 231–243.

(34) Wei, J.; Li, Y. CRISPR-Based Gene Editing Technology and Its Application in Microbial Engineering. Eng. Microbiol. 2023, 3, 100101–100103.

(35) Bloudoff, K.; Fage, C. D.; Marahiel, M. A.; Schmeing, T. M. Structural and Mutational Analysis of the Nonribosomal Peptide Synthetase Heterocyclization Domain Provides Insight into Catalysis. Proc. Natl. Acad. Sci. 2017, 114, 95–100.

(36) Durairaj, P.; Li, S. Functional Expression and Regulation of Eukaryotic Cytochrome P450 Enzymes in Surrogate Microbial Cell Factories. Eng. Microbiol. 2022, 2, 100011–100028.

(37) Pfeifer, B. A.; Admiraal, S. J.; Gramajo, H.; Cane, D. E.; Khosla, C. Biosynthesis of Complex Polyketides in a Metabolically Engineered Strain of E. coli. Science 2001, 291, 1790–1792.

(38) Quadri, L. E.; Quadri, L. E.; Weinreb, P. H.; Lei, M.; Nakano, M. M.; Zuber, P.; Walsh, C. Characterization of Sfp, a Bacillus subtilis Phosphopantetheinyl Transferase for Peptidyl Carrier Protein Domains in Peptide Synthetases. J. Biol. Chem. 1998, 37, 1585–1595.

(39) Omura, T.; Sato, R. The Carbon Monoxide-binding Pigment of Liver Microsomes. I. Evidence for Its Hemoprotein Nature. J. Biol. Chem. 1964, 239, 2370–2378.

(40) Zhang, W.; Du, L.; Qu, Z.; Zhang, X.; Li, F.; Li, Z.; Qi, F.; Wang, X.; Jiang, Y.; Men, P.; Sun, J.; Cao, S.; Geng, C.; Qi, F.; Wan, X.; Liu, C.; Li, S. Compartmentalized Biosynthesis of Mycophenolic Acid. Proc. Natl. Acad. Sci. 2019, 116, 13305–13310.

(41) Zhang, W.; Du, L.; Li, F.; Zhang, X.; Qu, Z.; Han, L.; Li, Z.; Sun, J.; Qi, F.; Yao, Q.; Sun, Y.; Geng, C.; Li, S. Mechanistic Insights into Interactions between Bacterial Class I P450 Enzymes and Redox Partners. ACS Catal. 2018, 8, 9992–10003.

(42) Kinsland, C.; Taylor, S. V.; Kelleher, N. L.; McLafferty, F. W.; Begley, T. P. Overexpression of Recombinant Proteins with a C-Terminal Thiocarboxylate: Implications for Protein Semisynthesis and Thiamin Biosynthesis. Protein Sci. 1998, 7, 1839–1842.

(43) Zheng, S.; Guo, J.; Cheng, F.; Gao, Z.; Du, L.; Meng, C.; Li, S.; Zhang, X. Cytochrome P450s In Algae: Bioactive Natural Product Biosynthesis and Light-Driven Bioproduction. Acta Pharm. Sin. B. 2022, 12, 1986–2001.

(44) Zhang, X. W.; Guo, J. W.; Cheng, F. Y.; Li, S. Y. Cytochrome P450 Enzymes in Fungal Natural Product Biosynthesis. Nat. Prod. Rep. 2021, 38, 1072–1099.

(45) Zhang, X.; Li, S. Expansion of Chemical Space for Natural Products by Uncommon P450 Reactions. Nat. Prod. Rep. 2017, 34, 1061–1089.

(46) Ushimaru, R.; Abe, I. C–N and C–S Bond Formation by Cytochrome P450 Enzymes. Trends Chem. 2023, 5, 526–536.

(47) Zocher, G.; Richter, M. E.; Mueller, U.; Hertweck, C. Structural fine-tuning of a multifunctional cytochrome P450 monooxygenase. J. Am. Chem. Soc. 2011, 133, 2292–2302.

(48) Guengerich, F. P. Kinetic Deuterium Isotope Effects in Cytochrome P450 Reactions. Methods Enzymol. 2017, 596, 217–238.

(49) Abramson, J.; Adler, J.; Dunger, J.; Evans, R.; Green, T.; Pritzel, A.; Ronneberger, O.; Willmore, L.; Ballard, A. J.; Bambrick, J. Nature, Accurate structure prediction of biomolecular interactions with AlphaFold 3. 2024, 630, 493–500.

(50) Trott, O.; Olson, A. AutoDock Vina: Improving the Speed and Accuracy of Docking with a New Scoring Function, Efficient Optimization, and Multithreading. J. Comput. Chem. 2010, 31, 455–461.

(51) Denisov, I. G.; Makris, T. M.; Sligar, S. G.; Schlichting, I. J. Structure and chemistry of cytochrome P450. Chem. Rev. 2005, 105, 2253–2278.

(52) Clark, J. P.; Miles, C. S.; Mowat, C. G.; Walkinshaw, M. D.; Reid, G. A.; Daff, S. N.; Chapman, S. K. The role of Thr268 and Phe393 in cytochrome P450 BM3. J. Inorg. Biochem. 2006, 100, 1075–1090.

(53) Kimata, Y.; Shimada, H.; Hirose, T.; Ishimura, Y. Role of Thr-252 in cytochrome P450cam: a study with unnatural amino acid mutagenesis. Biochem. Biophys. Res. Commun. 1995, 208, 96–102.

(54) Kuper, J.; Wong, T. S.; Roccatano, D.; Wilmanns, M.; Schwaneberg, U. J. Understanding a mechanism of organic cosolvent inactivation in heme monooxygenase P450 BM-3. J. Am. Chem. Soc. 2007, 129, 5786–5787.

(55) Kimata, Y.; Shimada, H.; Hirose, T.; Ishimura, Y. Role of Thr-252 in cytochrome P450cam: a study with unnatural amino acid mutagenesis. Biochem. Biophys. Res. Commun. 1995, 208, 96–102.

(56) Kuper, J.; Wong T, S.; Roccatano, D. Understanding a mechanism of organic cosolvent inactivation in heme monooxygenase P450 BM-3. J. Am. Chem. Soc. 2007, 129, 5786–5787.

(57) Belecki, K.; Townsend, C. A. Biochemical determination of enzyme-bound metabolites: preferential accumulation of a programmed octaketide on the Enediyne polyketide synthase CalE8. J. Am. Chem. Soc. 2013, 135, 14339–14348.

(58) Scharf, D. H.; Chankhamjon, P.; Scherlach, K.; Heinekamp, T.; Roth, M.; Brakhage, A. A.; Hertweck, C. Epidithiol formation by an unprecedented twin carbon-sulfur lyase in the gliotoxin pathway. Angew. Chem., Int. Ed. 2012, 51, 10064–8.

(59) Mihara, H.; Esaki, N., Bacterial cysteine desulfurases: their function and mechanisms. Appl. Microbiol. Biotechnol. 2002, 60, 12–23.

(60) Gaudelli, N. M.; Long, D. H.; Townsend, C. A. β-Lactam formation by a non-ribosomal peptide synthetase during antibiotic biosynthesis. Nature 2015, 520, 383–387.

(61) Wheadon, M. J.; Townsend, C. A. Evolutionary and functional analysis of an NRPS condensation domain integrates β-lactam, ?-amino acid, and dehydroamino acid synthesis. Proc. Natl. Acad. Sci. 2021, 118, e2026017118.

(62) Xue, Y.; Wang, X.; Liu, W. Reconstitution of the Linaridin Pathway Provides Access to the Family-Determining Activity of Two Membrane-Associated Proteins in the Formation of Structurally Underestimated Cypemycin. J. Am. Chem. Soc. 2023, 145, 7040–7047.

(63) Guarneros, G.; Ortega, M. V. Cysteine desulfhydrase activities of Salmonella typhimurium and Escherichia coli. Biochim. Biophys. Acta, Enzymol. 1970, 198 (1), 132–142.

(64) Higashi, T.; Ichikawa, T.; Inagaki, S.; Min, J. Z.; Fukushima, T.; Toyo’oka, T. Simple and practical derivatization procedure for enhanced detection of carboxylic acids in liquid chromatography– electrospray ionization-tandem mass spectrometry. J. Pharmaceut. Biomed. 2010, 52, 809–818.

(65) Pistofidis, A.; Ma, P.; Li, Z.; Munro, K.; Houk, K. N.; Schmeing, T. M. Structures and mechanism of condensation in non-ribosomal peptide synthesis. Nature 2024, 638 (8049), 270–278.

(66) Wu, P.; Yan, S.; Fang, W.; Wang, B. Molecular Mechanism of the Mononuclear Copper Complex-Catalyzed Water Oxidation from Cluster-Continuum Model Calculations. ChemSusChem 2022, 15, e202102508.

(67) Wang, Y.; Wang, L. Q.; Wu, F.; Liu, F.; Wang, Q.; Zhang, X. L.; Selvaraj, J. N.; Zhao, Y. J.; Xing, F. G.; Yin, W. B.; Liu, Y. A Consensus Ochratoxin A Biosynthetic Pathway: Insights from the Genome Sequence of Aspergillus ochraceus and a Comparative Genomic Analysis. Appl. Environ. Microb. 2018, 84 (19), e01009–18.

(68) Wang, B.; Cao, Z. Hydration of Carbonyl Groups: The Labile H3O+ Ion as an Intermediate Modulated by the Surrounding Water Molecules. Angew. Chem., Int. Ed. 2011, 50, 3266–3270.

(69) Dowling, D. P.; Kung, Y.; Croft, A. K.; Taghizadeh, K.; Kelly, W. L.; Walsh, C. T.; Drennan, C. L., Structural elements of an NRPS cyclization domain and its intermodule docking domain. Proc. Natl. Acad. Sci. 2016, 113 (44), 12432–12437.

(70) Gnann, A. D.; Xia, Y.; Soule, J.; Barthélemy, C.; Mawani, J. S.; Musoke, S. N.; Castellano, B. M.; Brignole, E. J.; Frueh, D. P.; Dowling, D. P. High-resolution structures of a siderophore-producing cyclization domain from Yersinia pestis offer a refined proposal of substrate binding. J. Bio. Chem. 2022, 298 (10), 102454.

(71) Pang, B.; Liao, R.; Tang, Z.; Guo, S.; Wu, Z.; Liu, W., Caerulomycin and collismycin antibiotics share a trans-acting flavoprotein-dependent assembly line for 2,2’-bipyridine formation. Nat. Commun. 2021, 12, 3124–3136.

(72) Van Cura, D.; Ng, T. L.; Huang, J.; Hager, H.; Hartwig, J. F.; Keasling, J. D.; Balskus, E. P. Discovery of the azaserine biosynthetic pathway uncovers a biological route for alpha-diazoester production. Angew. Chem., Int. Ed. 2023, e202304646.

(73) Chen, Y.; Tu, Y.; Pan, T.; Deng, Z.; Duan, L. A cysteine-reloading process initiating the biosynthesis of the bicyclic scaffold of dithiolopyrrolones. Antibiotics (Basel). 2023, 12, 787–799.

(74) Dong, L. B.; Rudolf, J. D.; Kang, D.; Wang, N.; He, C. Q.; Deng, Y.; Huang, Y.; Houk, K. N.; Duan, Y.; Shen, B. Biosynthesis of thiocar-boxylic acid-containing natural products. Nat. Commun. 2018, 9, 2362–2371.

(75) Notredame, C.; Higgins, D. G.; Heringa, J. T-Coffee: A novel method for fast and accurate multiple sequence alignment. J. Mol. Biol. 2000, 302, 205–217.

(76) Tamura, K.; Stecher, G.; Kumar, S. MEGA11: Molecular evolutionary genetics analysis version 11. Mol. Biol. Evol. 2021, 38, 3022–3027.

(77) Kieser, T.; Bibb, M. J.; Buttner, M. J.; Chater, K. F.; Hopwood, D. A. Practical Streptomyces Genetics. 2000, (John Innes Foundation Norwich,).

(78) Flett, F.; Mersinias, V.; Smith, C. P. High efficiency intergeneric conjugal transfer of plasmid DNA from Escherichia coli to methyl DNA-restricting Streptomycetes. FEMS Microbiol. Lett. 1997, 155, 223–229.

(79) Mossessova, E.; Lima, C. D. Ulp1-SUMO crystal structure and genetic analysis reveal conserved interactions and a regulatory element essential for cell growth in yeast. Mol. Cell. 2000, 5, 865–876.

(80) Sato, R. Cytochrome-P-450 - an inconspicuous start - a citation-classic commentary on the carbon monoxide-binding pigment of liver-microsomes .1. evidence for its hemoprotein nature. Cc/Life. Sci. 1991, 8, 9–9.

(81) Zhang, Y.; Liu, H.; Yang, S.; Luo, R.; Chen, H. Well-balanced force field ff 03 CMAP for folded and disordered proteins. J. Chem. Theory Comput. 2019, 15, 6769–6780.

(82) Zheng, L.; Meng, J.; Jiang, K.; Lan, H.; Wang, Z.; Lin, M.; Li, W.; Guo, H.; Wei, Y.; Mu, Y. Improving Protein–Ligand Docking and Screening Accuracies by Incorporating a Scoring Function Correction Term. Briefings Bioinf. 2022, 23, 1–15.

(83) Lu, Y.; Sen, K.; Yong, C.; Gunn, D. S. D.; Purton, J. A.; Guan, J.; Desmoutier, A.; Abdul Nasir, J.; Zhang, X.; Zhu, L.; Hou, Q.; Jackson- Masters, J.; Watts, S.; Hanson, R.; Thomas, H. N.; Jayawardena, O.; Logsdail, A. J.; Woodley, S. M.; Senn, H. M.; Sherwood, P.; Catlow, C. R. A.; Sokol, A. A.; Keal, T. W. Multiscale QM/MM modelling of catalytic systems with ChemShell. Phys. Chem. Chem. Phys. 2023, 25, 21816–21835.

(84) Metz, S.; Kästner, J.; Sokol, A. A.; Keal, T. W.; Sherwood, P. ChemShell—a modular software package for QM/MM simulations. WIREs Comput. Mol. Sci. 2014, 4, 101–110.

(85) Ahlrichs, R.; Bär, M.; Häser, M.; Horn, H.; Kölmel, C. Electronic structure calculations on workstation computers: The program system turbomole. Chem. Phys. Lett. 1989, 162, 165–169.

(86) Smith, W.; Forester, T. R. DL_POLY_2.0: A general-purpose parallel molecular dynamics simulation package. J. Mol. Graphics 1996, 14, 136–141.

(87) Smith, W.; Yong, C. W.; Rodger, P. M. DL_POLY: Application to molecular simulation. Mol. Simul. 2002, 28, 385–471.

(88) Bakowies, D.; Thiel, W. Hybrid models for combined quantum mechanical and molecular mechanical approaches. J. Chem. Phys 1996, 100, 10580–10594.

(89) Becke, A. D. Density-functional thermochemistry. III. The role of exact exchange. J. Chem. Phys 1993, 98 (7), 5648–5652.

(90) Schäfer, A.; Horn, H.; Ahlrichs, R. Fully optimized contracted Gaussian basis sets for atoms Li to Kr. J. Chem. Phys 1992, 97 (4), 2571–2577.

(91) Grimme S. Semiempirical GGA-Type density functional constructed with a Long-range dispersion correction. J. Comput. Chem. 2006, 27, 1787–1799.

(92) Grimme S.; Ehrlich S.; Goerigk L. Effect of the damping function in dispersion corrected density functional theory. J. Comput. Chem. 2011, 32, 1456–1465.

